# Cell confinement reveals a branched-actin independent circuit for neutrophil polarity

**DOI:** 10.1101/457119

**Authors:** Brian R. Graziano, Jason P. Town, Tamas L. Nagy, Miha Fošnarič, Samo Penič, Aleš Iglič, Veronika Kralj-Iglič, Nir Gov, Alba Diz-Muñoz, Orion D. Weiner

**Author notes:** To whom correspondence should be addressed: Orion Weiner, CVRI & Department of Biochemistry and Biophysics, University of California, San Francisco, Cardiovascular Research Building, Room 352F, MC 3120, 555 Mission Bay Blvd. South, San Francisco, CA 94158.

## Abstract

Migratory cells use distinct motility modes to navigate different microenvironments, but it is unclear whether these modes rely on the same core set of polarity components. To investigate this, we disrupted Arp2/3 and WAVE complex, which assemble branched actin networks that are essential for neutrophil polarity and motility in standard adherent conditions. Surprisingly, confinement rescues polarity and movement of neutrophils lacking these components, revealing a processive bleb-based protrusion program that is mechanistically distinct from the branched actin-based protrusion program but shares some of the same core components and underlying molecular logic. We further find that the restriction of protrusion growth to one site does not always respond to membrane tension directly, as previously thought, but may rely on closely linked properties such as local membrane curvature. Our work reveals a hidden circuit for neutrophil polarity and indicates that cells have distinct molecular mechanisms for polarization that dominate in different microenvironments.

## INTRODUCTION

Directed migration underlies a wide array of biological processes, including embryogenesis, wound healing, and the ability of immune cells to track and destroy pathogens. A key step in migration is polarization, where cells restrict the activity of their protrusive machinery to a limited portion of their surface. Polarization is thought to arise through the coordinated interactions of short-range positive and long-range negative feedback loops, with actin polymerization playing essential roles in both types of feedback (Houk et al., 2012; Inoue and Meyer, 2008; Keren et al., 2008; Meinhardt, 1999; Neilson et al., 2011; Nguyen et al., 2016; Sasaki et al., 2007; Weiner et al., 2007; Xiong et al., 2010).

When migratory cells first sense a spatially uniform increase in stimulatory cues, they respond by rapidly activating leading-edge polarity factors, including RhoGTPases (e.g., Rac), throughout the plasma membrane (Weiner et al., 2007; Yang et al., 2016). These GTPases, in turn, activate Arp2/3-driven assembly of branched actin networks by recruiting nucleation-promoting factors (NPFs) such as WAVE and WASP (Fritz-Laylin et al., 2017; Machesky and Insall, 1998; Veltman et al., 2012; Weiner et al., 2006), resulting in the formation of multiple sheet-like protrusions. Each of these nascent polarity sites is sustained by multiple short-range positive feedback loops (e.g., recruitment of additional RhoGTPase activators (Nguyen et al., 2016)), in a manner that often depends on actin assembly (Inoue and Meyer, 2008; Srinivasan et al., 2003; Weiner et al., 2002). In parallel with this initial step of activation—but occurring on a slower timescale—cells generate long-range negative feedback to enable a dominant front to emerge (Swaney et al., 2010). In neutrophils, cells of the innate immune system, this process occurs through the growth of actin filaments, which stretch and increase tension in the plasma membrane (Houk et al., 2012). This membrane tension is thought to act as a long-range inhibitor of actin nucleation and polymerization, which enables a “winner-takes-all” competition among the nascent polarity sites to produce a single protrusion (Fig. 1A) (Diz-Muñoz et al., 2016; Houk et al., 2012; Keren et al., 2008; Sens and Plastino, 2015).

**Figure 1.**
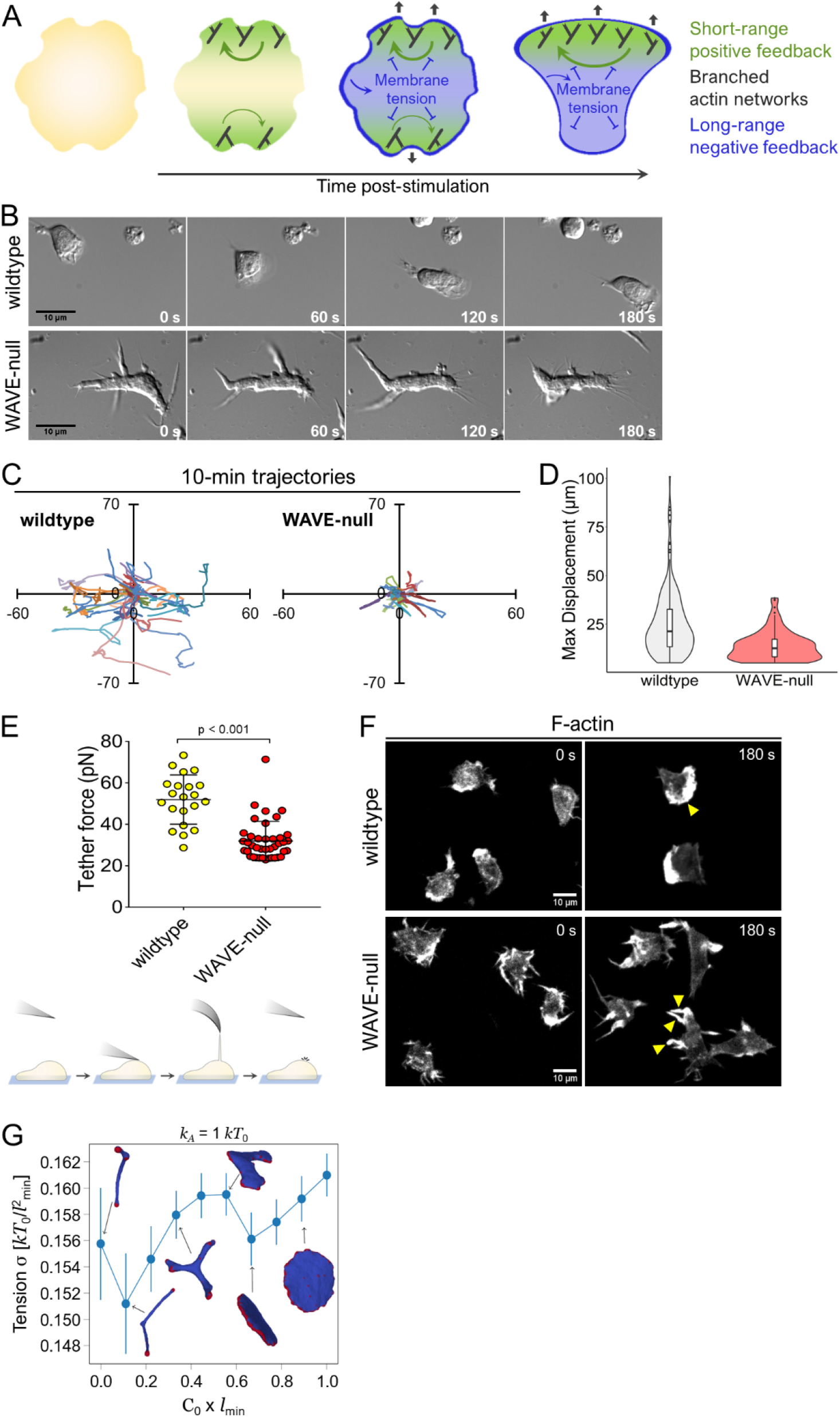
WAVE complex is required for neutrophil polarity and motility. **A)** Polarity generation depends on fast-acting, short-range positive feedback and slower-acting, long-range negative feedback. Actin assembly participates in both types of feedback, with its role in negative feedback arising from mechanical force production at the plasma membrane. **B)** Migration of differentiated HL-60 cells (dHL-60s) on fibronectin-coated glass in uniform chemoattractant (10 nM fMLP) visualized with differential interference contrast (DIC) microscopy. **C-D)** dHL-60s were prepared as in B. Cells were tracked using NucBlue nuclear stain and imaged every 15 s by widefield epifluorescence microscopy. **C)** Representative tracks from 30 wildtype and 30 WAVE-null cells (randomly selected) over a 10-min observation window. Axes, distance in μm. **D)** Maximum displacement measurements. Each point in the distribution represents the furthest a cell moved relative to its initial starting position over the 10-min observation period. N = 338 wildtype and 278 WAVE-null cells pooled from 2 independent experiments. **E)** Top, Membrane tether force was measured using atomic force microscopy (AFM) in dHL-60s cells plated for at least 10 min in 10 nM fMLP. Each point represents a single cell. Data were pooled from 3 independent experiments. Error bars, SD. Bottom, Schematic depicting the experimental procedure. An AFM cantilever was briefly brought into contact with a cell, then withdrawn. In instances when a membrane tether was successfully pulled, the cantilever was maintained at a constant height until the tether broke. The change in cantilever deflection was used to measure the ‘tether force’. **F)** dHL-60s in suspension were fixed and stained for F-actin using rhodamine-phalloidin in the presence or absence of chemoattractant (100 nM fMLP). Yellow arrowheads, actin-rich protrusions. **G)** Computational simulations calculating membrane tension σ (see “Materials and Methods”) as a function of spontaneous curvature, c0, of actin nucleators. Averaging was performed over 200 statistically independent microstates in equilibrium. Actin nucleators, red. Protein-free lipid bilayer, blue. Error bars, SD.

While actin polymerization serves as a key ingredient in generating the positive and negative feedback loops that give rise to polarity, we lack an understanding of how specific types of actin networks provide each kind of feedback. Immune cells assemble multiple actin networks at different subcellular locations that carry out distinct functions to support migration: Arp2/3-dependent assembly of branched actin networks at the leading edge contributes to cell guidance/steering and protrusion extension, while actomyosin bundles near the trailing edge provide contractile force to lift the cell rear and squeeze the cell body forward (Lämmermann and Germain, 2014; Moreau et al., 2018). Along with these functional differences, the types of actin networks immune cells and other migratory cells employ for migration vary with microenvironment (Lämmermann and Germain, 2014; Paluch et al., 2016). The role of actin dynamics in migration is complex and likely depends on the type of actin network, its subcellular location, and the extracellular environment. Existing tools to probe the role of actin networks in both the positive and negative feedback loops needed for polarity are fairly crude and have largely been based on pharmacological perturbations that target all actin polymer (Diz-Muñoz et al., 2016; Huang et al., 2013; Inoue and Meyer, 2008; Sasaki et al., 2007; Wang et al., 2002; Weiner et al., 2007; Yang et al., 2016). More surgical experiments are needed to clarify how different subcellular actin networks contribute to polarity generation under different environmental conditions.

Here, we address this question by dissecting the role of branched actin assembly in the neutrophil polarity program. We use CRISPR-mediated genome editing in human neutrophils (differentiated HL-60 cells) to knock out either the Arp2/3 complex, the nucleator of branched actin assembly (Mullins et al., 1998; Welch et al., 1997), or its key activator the WAVE complex (Machesky et al., 1999), which promotes branched actin assembly at the leading edge of migratory cells (Davidson et al., 2018; Kurisu et al., 2005; Leithner et al., 2016; Weiner et al., 2006; Yamazaki et al., 2003). We find that during unconfined migration, Arp2/3 and WAVE complex are each required for the mechanical force generation that supports long-range negative feedback and restriction of protrusion growth to one location. However, each complex is dispensable for polarity and movement of confined cells, where cell-*extrinsic* mechanical forces can compensate for the cell-*intrinsic* forces normally produced by WAVE complex. This confined movement relies on a completely different mode of leading-edge advance, with processive bleb-based protrusions forming in a back-and-forth motion that extends the leading edge in a serpentine manner. Surprisingly, these serpentine protrusions coincide with Rac-based local positive feedback loops that set a permissive zone for bleb propagation, operate independently of branched actin assembly, and form a mechanistically distinct polarity circuit in this context. Finally, we find that the long-range inhibitor that underlies competition between protrusions does not always respond to membrane tension directly but may rely on closely linked properties such as global cell shape and local membrane curvature. Our data indicate that multiple mechanistically distinct, but conceptually similar, polarity circuits operate in different cell migration contexts.

## RESULTS

### WAVE complex is required for polarization during adhesion-dependent migration

The WAVE complex is required for proper leading edge formation in immune cells (Leithner et al., 2016; Weiner et al., 2006) and its role in regulating cell shape and motility is broadly conserved across metazoans (Blagg et al., 2003; Kunda et al., 2003; Shakir et al., 2008). To generate a human neutrophil cell line lacking the WAVE complex (hereafter referred to as ‘WAVE-null’), we used CRISPR-Cas9 to knock out *HEM1/NCKAPL1*, which encodes the sole hematopoietic-specific component of the pentameric WAVE complex (Weiner et al., 2006), in dHL-60 cells. Disruption of *HEM1* is preferable to targeting *WAVE2/WASF2* directly since loss of HEM1 results in the concomitant degradation of the other remaining subunits of WAVE complex, including WAVE2 (Fig. S1A), and avoids potentially confounding gain-of-function effects of partial sub-complexes (Leithner et al., 2016; Litschko et al., 2017; Weiner et al., 2006).

As expected, WAVE-null cells showed severe polarity and morphological defects when plated in a standard 2-D adhesion setting. Treatment of WAVE-null cells with uniform chemoattractant resulted in the production of multiple dynamic finger-like protrusions radiating from the cell surface in a non-polarized fashion, rather than a single smooth leading edge typically observed in wildtype cells (Fig. 1B **and Videos 1-2**). WAVE-null cells also showed a pronounced motility defect during chemokinesis: When observed over a 10-min period, WAVE-null cells showed a mean maximum displacement of 14 ± 1 µm from their initial positions vs. 25 ± 1 µm for wildtype cells (Fig. 1C-D). The phenotypes we observed are broadly similar to those reported in mouse dendritic cells lacking WAVE complex (Leithner et al., 2016), in that both cell types failed to form lamellipod-like protrusions at the leading edge. However, ‘WAVE-null’ mouse dendritic cells showed only mildly impaired motility and appeared to have only a minor polarity defect. These differences may arise from the distinct physiological functions these two cell types perform (Lämmermann and Germain, 2014). We also note that the polarity defects we observed in WAVE-null neutrophils were more severe than those reported in prior work using RNAi to knockdown the HEM1 subunit of WAVE complex (Weiner et al., 2006). As our CRISPR-based approach eliminates all protein, the milder defects observed upon HEM1 knockdown may arise from residual WAVE complex following RNAi.

The multipolar phenotype we observed in WAVE-null neutrophils could arise from a defect in membrane tension generation, which may be needed for producing the negative feedback that fronts use to compete with one another (Houk et al., 2012; Raucher and Sheetz, 2000) (Fig. 1A). To test this idea, we used atomic force microscopy (AFM) to measure membrane tether force in cells stimulated with chemoattractant (Fig. 1E). WAVE-null cells showed a two-fold decrease in tether force vs. wildtype cells (Fig. 1E), corresponding to a four-fold decrease in membrane tension when using a common model relating tether force to membrane tension (Diz-Muñoz et al., 2016; Houk et al., 2012); see (Hochmuth et al., 1996) for calculation details. These decreases were observed despite WAVE-null cells forming multiple actin-rich protrusions in response to stimulation with chemoattractant (Fig. 1F). Our results show that WAVE-dependent actin assembly is required for efficient generation of membrane tension increases.

We further performed computational simulations to assess whether changes in the spatial organization of actin nucleation can correlate with changes in membrane tension. By modulating the curvature preference of a fixed concentration of actin nucleators, we generated membrane surfaces ranging from those containing multiple finger-like protrusions (similar to WAVE-null cells) to a single sheet-like protrusion (similar to wildtype cells). Transition in membrane shape from finger-like to sheet-like coincided with small increases in membrane tension (Fig. 1G **and** S2). These simulations show that even if the concentration of actin nucleators remains constant, changes in their organization can strongly affect the morphology of the resulting protrusions they form and, consequently, the amount of membrane tension they can generate. In combination with our experimental data, they suggest that spatial organization of WAVE complex at the plasma membrane is a major factor contributing to WAVE complex’s ability to generate lamellipod-like protrusions and that protrusions with this morphology may be more effective at increasing membrane tension (and membrane tension-linked cell processes like membrane geometry) than other protrusion types.

**Figure 2.**
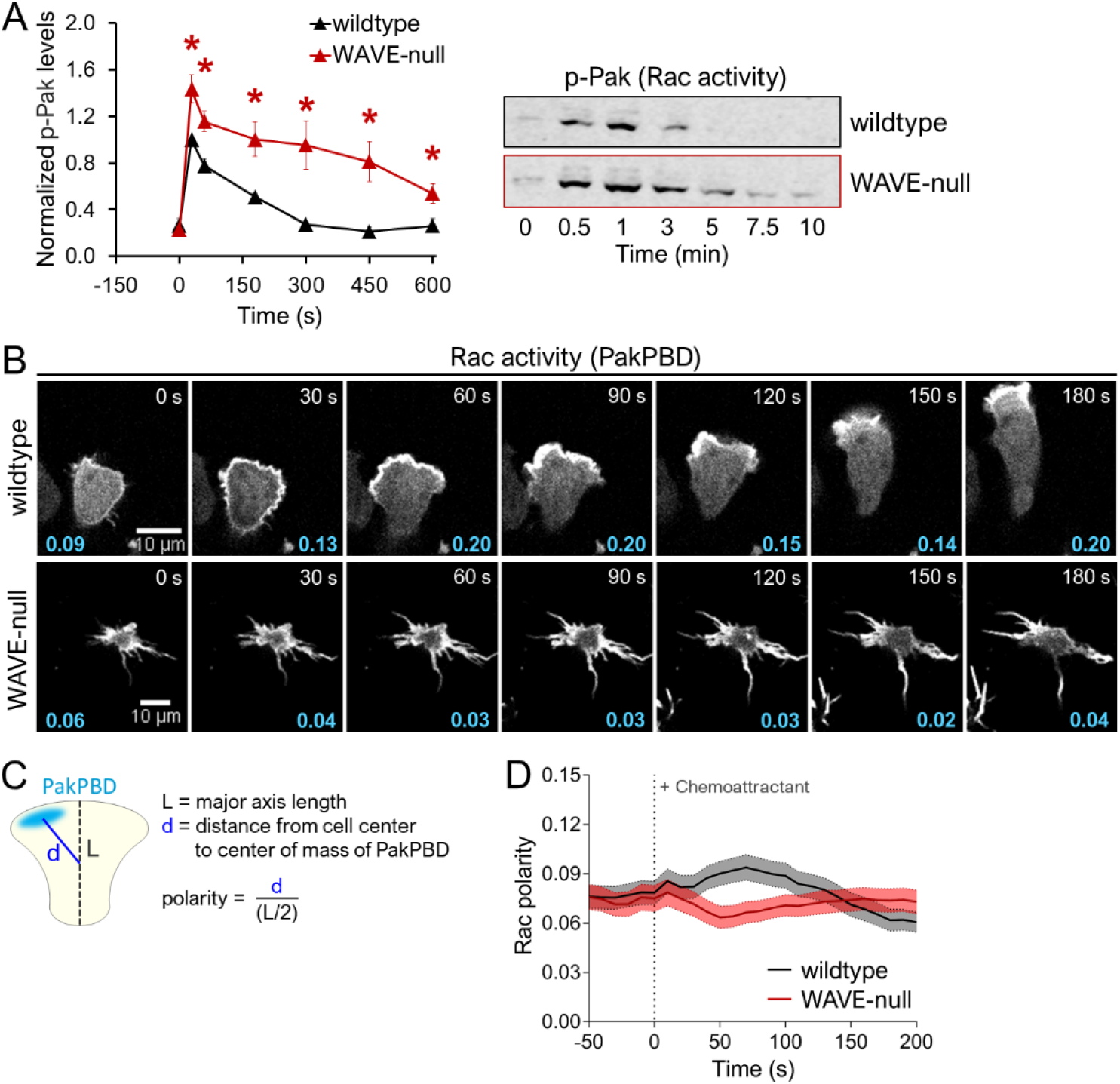
Loss of WAVE complex results in aberrant levels and polarization of Rac activity. **A)** Rac activity was quantified for chemoattractant-stimulated cells using antibodies targeting phospho-Pak, a downstream readout of Rac activation. Antibodies targeting total Pak were used as loading controls. Left, each point represents an average of 4 independent experiments, with data for each experiment normalized to wildtype cells at ‘0 s’. Error bars, SEM. *, P < 0.05 by unpaired t-test. Right, representative immunoblot. **B)** dHL-60 cells expressing the Rac biosensor PakPBD-mCitrine were plated on fibronectin-coated glass, stimulated with 100 nM fMLP, and imaged by confocal microscopy every 10 s. Values in cyan indicate the degree of PakPBD polarization, as described in D. **C)** Schematic depicting calculation of Rac activity polarity. See “Materials and Methods” for details. **D)** dHL-60s were prepared as in B, and polarity of Rac activity was measured for single cells at each 10-s interval. Dark lines, mean polarity of Rac activity. Light shaded regions, +/− 95% CI of the mean. N = 211 wildtype and 198 WAVE-null cells pooled from 2 independent experiments.

### Key leading-edge polarity factors require Arp2/3 and WAVE complex for proper regulation

Membrane tension is thought to play a major role in negatively regulating the polarity factors that organize protrusion growth, including the RhoGTPase Rac (Houk et al., 2012; Lieber et al., 2013). The morphological polarity defects observed in WAVE-null cells are expected based on the decreased membrane tension in these cells, which should fail to engage the global negative feedback circuit that constrains the amounts and spatial distribution of Rac activity. We tested this hypothesis by monitoring the levels of Rac activation using the phosphorylation state of Pak kinase, a Rac effector (Knaus and Bokoch, 1998). Following stimulation, WAVE-null cells produced significantly higher levels of phospho-Pak than wildtype cells (Fig. 2A), consistent with elevated levels of Rac activity. This result is also in agreement with recent work revealing a role for actin polymerization in negatively regulating Rac activity in neutrophils (Graziano et al., 2017).

We next examined whether elevated levels of Rac activity coincided with failure to restrict Rac activity to a single site. Using the PakPBD biosensor (Manser et al., 1994; Weiner et al., 2007), we monitored the spatial distribution of Rac activity in live cells. Stimulation of wildtype cells with uniform chemoattractant produced an initial burst of Rac activity throughout the plasma membrane, followed by its restriction to a single protrusion (Fig. 2B, top row; **Video 3**), in agreement with prior work (Weiner et al., 2007; Yang et al., 2016). WAVE-null cells, in contrast, maintained multiple protrusions enriched for Rac activity for the entire timecourse (Fig. 2B, bottom row; **Video 4**). We quantified polarity in these cells by measuring the distance between the geometric cell center and the center of weighted Rac activity signal, followed by normalization using the cell length (Fig. 2C). This yields a ‘Rac polarity’ score from 0 to 1, with a value of ‘0’ indicating Rac activity distributed uniformly around the cell and a value near ‘1’ indicating a single focus of Rac activity on the portion of the membrane furthest from the cell center (see Methods for details). Populations of wildtype neutrophils polarized Rac activity in response to uniform chemoattractant, with cells reaching peak polarity around 60 s (Fig. 2D). In contrast, WAVE-null cells showed no Rac polarity increase following stimulation with chemoattractant (Fig. 2D), consistent with impaired spatial regulation of Rac activity.

A major cellular function of the WAVE complex is stimulation of Arp2/3-dependent branched actin assembly, but the WAVE complex’s ability to regulate Rac could also stem from its additional roles. In other systems, the WAVE complex interacts with regulators of Rac (e.g., Rac GAPs (Soderling et al., 2002)) or additional actin assembly factors (e.g., formins (Beli et al., 2008)). Furthermore, the WAVE complex may contribute to the growth of actin networks nucleated by WAVE-*independent* processes (Bieling et al., 2018). To test whether such Arp2/3-*independent* functions of the WAVE complex may contribute to its role in Rac regulation, we performed additional experiments in cells lacking functional Arp2/3 complex (Fig. S1B). These ‘ARP2-null’ cells phenocopied the key defects we observed in WAVE-null neutrophils: ARP2-null cells showed impaired membrane tension generation (Fig. 3A), elevated levels of Rac activity (Fig. 3B), and diminished polarization of Rac activity in response to stimulation with uniform chemoattractant (Fig. 3C-D). These data are consistent with WAVE complex primarily regulating polarity through its stimulation of branched actin assembly rather than through its other cellular functions.

**Figure 3.**
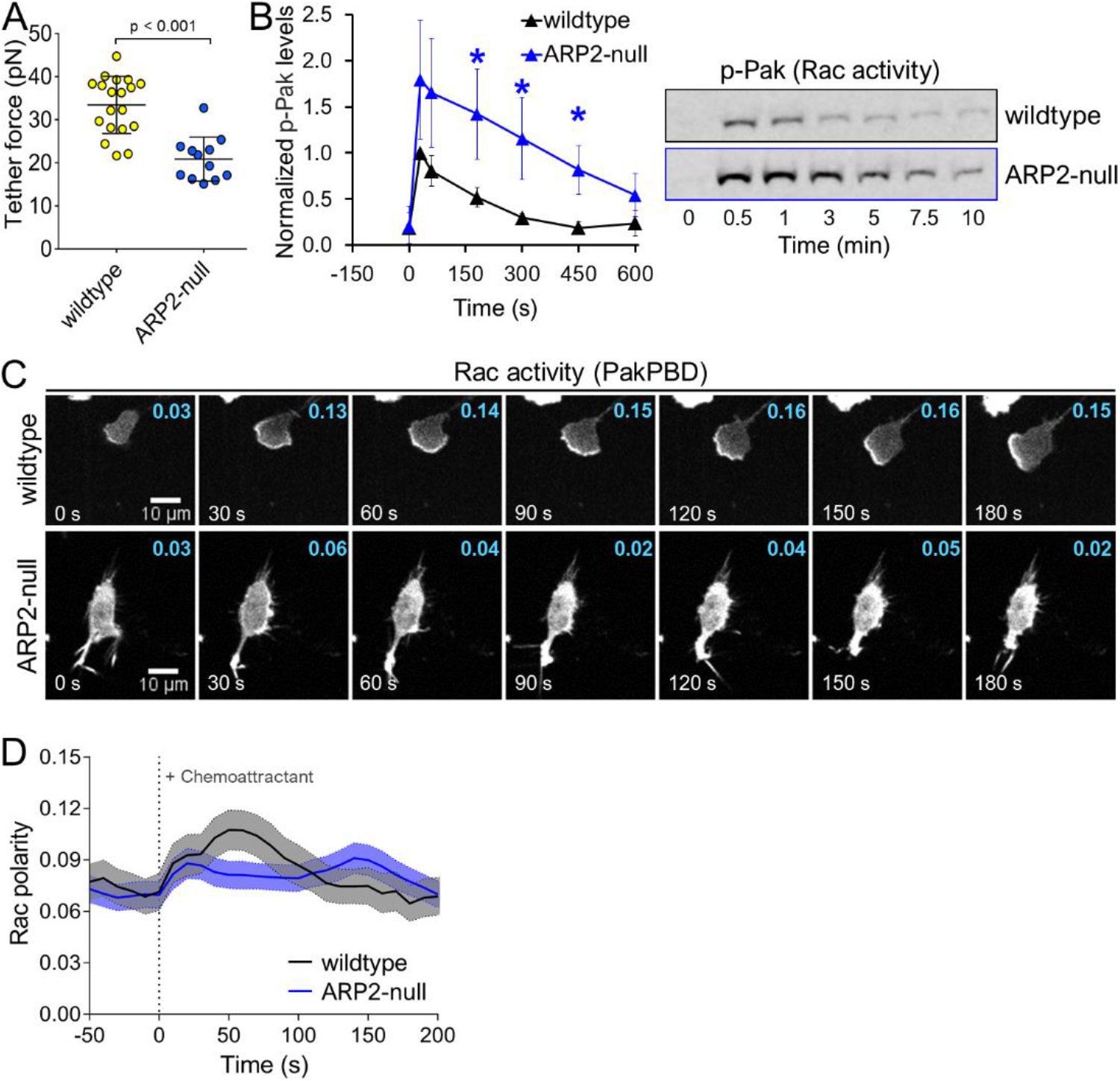
Loss of Arp2/3 complex phenocopies polarity defects observed in WAVE-null cells. **A)** Membrane tether force of dHL-60s was measured using AFM as described in 1E. Each point represents a single cell. Data were pooled from 2 independent experiments. Error bars, SD. **B)** dHL-60s were stimulated with 10 nM fMLP and samples were collected and processed for immunoblot. Left, Rac activity was indirectly quantified as in 2A. Each point represents an average of 4 independent experiments, with data for each experiment normalized to wildtype cells at ‘0 s’. Error bars, SEM. *, P < 0.05 by unpaired t-test. Right, representative immunoblot. **C)** dHL-60 cells expressing PakPBD-mCitrine were imaged as in 2B. Values in cyan indicate the degree of PakPBD polarization, as described in 2C. **D)** Quantification of Rac polarity as described in 2C for cells prepared as in 3C. Dark lines, mean Rac polarity. Light shaded regions, +/− 95% CI of the mean. N = 101 wildtype and 185 ARP2-null cells pooled from 2 independent experiments.

### Hypotonic treatment restores polarity in the absence of WAVE complex

If the WAVE complex primarily regulates long-range inhibition of leading-edge signals by imparting mechanical force on the plasma membrane, we should be able to restore long-range inhibition by providing compensatory mechanical forces. One approach for mimicking WAVE’s extension of the plasma membrane is to place neutrophils in hypotonic media, where osmotic-based cell swelling may provide such a force (Fig. 4A) (Gauthier et al., 2011; Houk et al., 2012). Since wildtype neutrophils can still polarize and migrate in 0.5x isotonic media (i.e. a 1:1 dilution of isotonic media with deionized water) (Diz-Muñoz et al., 2016), we examined how stimulating WAVE-null cells under similar conditions affected the amount of Rac activity. In hypotonic media (0.5x isotonic), both wildtype and WAVE-null cells showed nearly identical levels of Rac activity throughout the entire timecourse (as assayed via Pak phosphorylation, Fig. 4B), indicating a rescue of the Rac activity defect observed in WAVE null cells in isotonic media (Fig. 2A). These data suggest that hypotonic treatment can supply the mechanical force that is normally provided by WAVE-dependent actin assembly to generate the negative feedback needed for limiting total cellular Rac activity.

**Figure 4.**
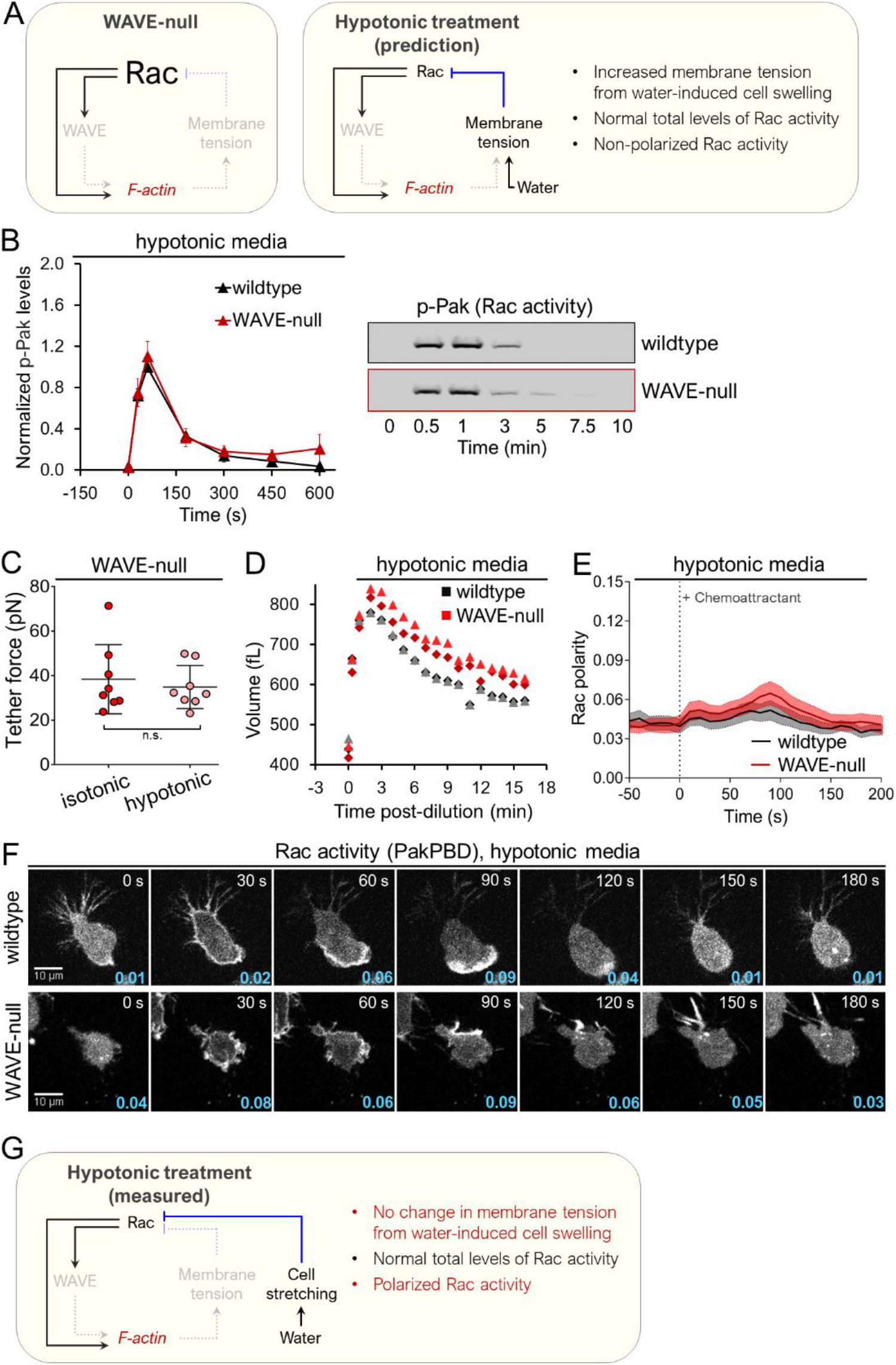
Hypotonic treatment to extend the plasma membrane restores spatiotemporal Rac activity in cells lacking WAVE complex. **A)** Left, schematic depicting key components in the neutrophil polarity circuit and their measured alterations in WAVE-null cells. Right, expected regulatory changes among key polarity components in WAVE-null cells following hypotonic-induced plasma membrane stretching, where we hypothesized a rescue of Rac activity levels. **B)** dHL-60s were suspended in 0.5x isotonic media (see “Materials and methods” for details), stimulated with 10 nM fMLP and samples were collected and processed for immunoblot. Left, Rac activity was quantified via phospho-Pak as in 3B. Each point represents an average of 4 independent experiments, with data for each experiment normalized to wildtype cells at ‘0 s’. Error bars, SEM. Right, representative immunoblot. **C)** Membrane tether force of dHL-60s was measured using AFM as described in 1E. The same cells were measured before and after hypotonic shock. Note that the cells in the ‘isotonic’ condition are a subset of the data depicted in 1E that were further subjected to hypotonic treatment. The data are reproduced here to aid comparison. Each point represents a single cell. Data were pooled from 2 independent experiments. Error bars, SD. Membrane tension does not significantly change following hypotonic shock. **D)** Volumes of dHL-60 cells in suspension following hypotonic treatment (0.5x isotonic media) were measured using a Coulter Counter. Each point represents the mode volume for a distribution of >7000 cells (see “Materials and Methods” for details). Data are pooled from 2 independent experiments. Volume significantly increases following hypotonic shock. **E)** Quantification of Rac polarity as described in 2C for cells prepared as in 4F. Dark lines, mean polarity of Rac activity. Light shaded regions, +/− 95% CI of the mean. N = 148 wildtype and 129 WAVE-null cells pooled from 2 independent experiments. **F)** dHL-60 cells in hypotonic media (0.5x isotonic) expressing the Rac activity reporter PakPBD-mCitrine were imaged as in 2B. Values in cyan indicate the degree of Rac polarization, as described in 2C. **G)** Schematic depicting measured regulatory changes among key polarity components in WAVE-null cells following hypotonic treatment. Compare with expected changes in 4A.

We next examined what physical changes the WAVE complex might impart on the plasma membrane to regulate Rac activity. Prior work concluded that membrane tension increases play a primary role in this regulatory behavior: Experimental perturbations that increase membrane tension lead to inhibition of both Rac activity and the WAVE complex; conversely, decreases in membrane tension lead to their activation throughout the cell (Houk et al., 2012). However, tension increases frequently coincide with other physical changes (e.g., unfolding of membrane reservoirs (Hallett et al., 2008; Sens and Plastino, 2015)), and it is not clear whether membrane tension itself or a close correlate forms the basis of the long-range inhibition. Determining whether membrane tension or other physical parameters execute the long-range inhibition is critical to identifying the relevant mechanosensor that orchestrates this process (Diz-Muñoz et al., 2013). Using AFM to measure membrane tether force, we were surprised to find that mild hypotonic treatment of WAVE-null cells did not increase membrane tension (Fig. 4C), so the rescue of Rac activity by hypotonic treatment is not a consequence of increased membrane tension. However, hypotonic shock did cause other alterations to the organization of the membrane of WAVE-null cells that normally go along with tension changes. In particular, volume increased by as much as two-fold (Fig. 4D), indicating a significant increase in the apparent surface area of the membrane (Hallett et al., 2008). These observations are consistent with leukocytes maintaining membrane reservoirs that can be released in response to osmolarity decreases (Cheung et al., 1982; Ting-Beall et al., 1993). Furthermore, they indicate that polarity-regulating mechanosensors respond to physical changes in the plasma membrane beyond tension increases.

As hypotonic treatment resulted in both wildtype and WAVE-null cells producing similar amounts of total Rac activity, we assessed whether this perturbation might also restore polarization of Rac activity in WAVE-null cells using quantification of PakPBD localization at previously described (Fig. 2C). We found that WAVE-null cells in 0.5x isotonic media could mildly polarize Rac activity in response to chemoattractant (Fig. 4E-F; **Videos 5-6**), whereas they had failed to do so in isotonic media (Fig. 2B **and** 2D). However, hypotonic treatment did not affect protrusion morphology, which remained finger-like (Fig. 4F, bottom row). Our results show that other mechanisms may function independent of the WAVE complex to polarize Rac activity, so long as the requirement for long-range inhibition (i.e. mechanical force) is satisfied—in this case by hypotonic-induced cell swelling (Fig. 4G).

### Arp2/3 and WAVE complex are dispensable for polarization in confined environments

Since hypotonic treatment has the potential to introduce pleiotropic effects beyond cell shape changes, we employed an orthogonal approach to stretch cells using mechanical force. Using polydimethylsiloxane (PDMS)-based devices, we created confined environments where chamber height could be adjusted by applying vacuum (Le Berre et al., 2012). We initially confined neutrophils by adjusting the height of the chamber to <5 μm. Acute stimulation of cells in this device was achieved using ultraviolet light to release chemically-caged chemoattractant (Collins et al., 2015; Pirrung et al., 2000) (Fig. 5A), and polarization of Rac activity was monitored using PakPBD. As expected, wildtype cells under confinement produced chemoattractant-based increases in polarized Rac activity (Fig. 5B-C; **Videos 7-8**). These responses broadly resembled those we observed in wildtype cells during unconfined migration (Fig. 2B **and** 2D).

**Figure 5.**
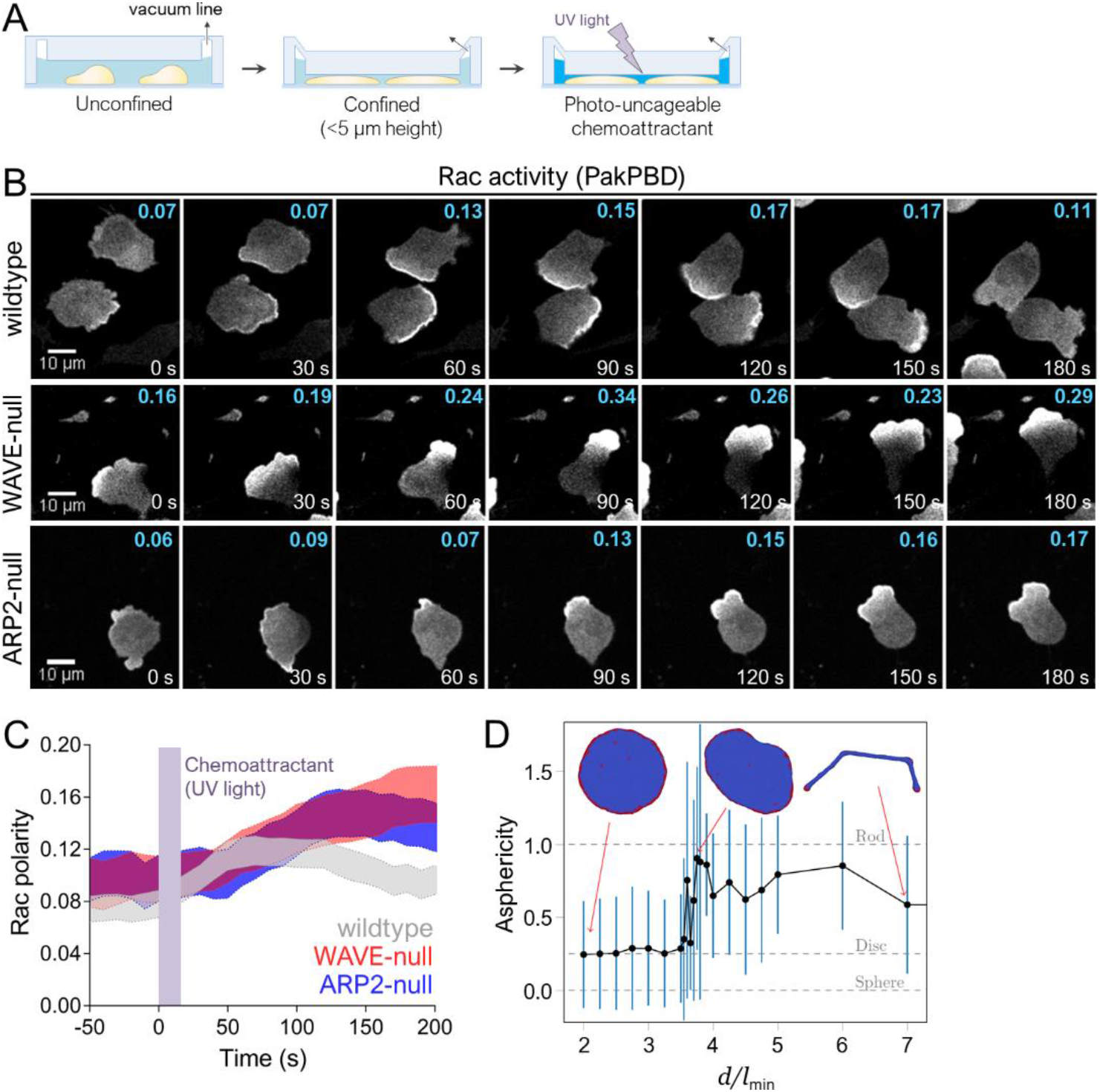
Cell compression restores polarization in cells lacking WAVE or Arp2/3 complex. **A)** Schematic for cell confinement experiments. The height of the chamber was set using a vacuum regulator. **B)** dHL-60 cells expressing the Rac biosensor PakPBD were plated on fibronectin-coated glass in media containing 10 μM caged-fMLP and imaged every 10 s by confocal microscopy. Chamber height was set as shown in 5A. Values in cyan indicate the degree of Rac activity polarization, as described in 2C, for the topmost cell fully inside each panel. **C)** Quantification of Rac polarity as described in 2C for cells prepared as in 5B. Violet bar indicates when UV was used to photo-uncage caged-fMLP (0-20 s). Shaded regions, +/− 95% CI of the mean Rac polarity. N = 134 wildtype cells pooled from 3 independent experiments; 73 WAVE-null cells pooled from 3 independent experiments; and 64 ARP2-null cells pooled from 2 independent experiments. **D)** Ensemble averaged asphericity as a function of distance d between two parallel plates, as determined from computational simulations. Asphericity is zero for sphere, 0.25 for very thin disc and 1 for very thin rod (gray dashed horizontal lines). Black dots indicate asphericity averaged over an ensemble of 500 statistically uncorrelated microstates and blue bars denote SD. *l_min_* is the edge length of the triangles in the mesh used to cover the surface of the vesicles. d is expressed in units of *l_min_*. Representative vesicle snapshots are shown for *d*/*l*_min_ = 2, 3.75 and 7. Actin nucleators denoted by red vertices, protein-free lipid bilayer by blue.

Whereas unconfined WAVE-null cells failed to polarize Rac activity when stimulated, confinement gave a profound rescue of polarity. Cells switched from producing finger-like protrusions to generating smooth leading edges that much more closely resembled the overall shape of wildtype cells (Fig. 5B, middle row; **Video 10**). These alterations in polarity and morphology were rapid and reversible. Repeatedly raising and lowering the height of the chamber (to alter the degree of confinement) caused WAVE-null neutrophils to switch from generating clustered finger-like protrusions to making smooth protrusions (**Video 11**). Stimulation of these confined WAVE-null neutrophils resulted in pronounced polarization of Rac activity that was even more profound than in wildtype cells under this degree of confinement (Fig. 5C). Experiments performed in ARP2-null cells produced similar results (Fig. 5B-C), further underscoring that all branched actin assembly is dispensable for supporting Rac polarity, leading edge morphology, and persistent motility.

We performed additional experiments under conditions where cells were only weakly confined (chamber height of 5-9 μm). While wildtype, WAVE-null, and ARP2-null cells all exhibited polarized Rac activity when stimulated with chemoattractant, we found that WAVE-null cells did not polarize Rac activity as strongly as wildtype or ARP2-null cells (Fig. S3A-C). Furthermore, WAVE-null cells at this confinement regime generated finger-like rather than smooth protrusions (Fig. S3B, middle row). Our observations in weakly confined WAVE-null cells are similar to those in hypotonically-treated WAVE-null cells (Fig. 4F, bottom row), where polarized Rac activity was rescued despite cells still forming protrusions of aberrant morphology. These results show that polarization of key leading-edge factors can be decoupled from protrusion morphology and may serve as a useful platform in future studies focusing on the mechanisms underlying each.

We then performed computational simulations to assess what physical processes might underlie the ability of WAVE-null cells to form smooth protrusions under compression. Using the system described in Fig. 1G (see also Materials and Methods), we observed how a fixed concentration of actin nucleators having zero (or small) spontaneous curvature (which produced vesicles with finger-like protrusions in an unconfined setting) affected vesicle morphology in response to varying degrees of compression. Surprisingly, we found that increasing levels of confinement (lower values of *d*/*l_min_*) resulted in vesicles converting from a morphology of primarily finger-like protrusions to one containing a single sheet-like protrusion (Fig. 5D **and Videos 12-14**). While this simplified model does not account for many other physical properties that neutrophils rely on to regulate shape and polarity (e.g., short range positive feedback, long range inhibition, myosin contractility), it does generally recapitulate our observations in WAVE-null cells where strong compression produced cells with broad smooth protrusions (Fig. 5B, middle row) but more weakly compressed cells retained finger-like protrusions (Fig. S3B, middle row). Furthermore, when WAVE-null cells are subjected to intermediate levels of compression (e.g., ceiling height = 5 µm), they alternate between forming sheet-like and finger-like protrusions (**Video 15**), similar to simulations of vesicles at intermediate levels of confinement (**Video 13**). Our simulations revealed that a key parameter specifying protrusion morphology is the ratio of actin nucleator protrusive force to the restoring force due to membrane bending, which determines the diameter of the long protrusions formed by the flat actin nucleators (see Supplementary Information and Fošnarič et al., 2018). When this value is lower than some critical value (i.e., under strong compression) the clusters of actin nucleators cannot form finger-like protrusions, and instead tend to spread uniformly along the free (unconfined) edge of the vesicle, which then takes the form of sheet-like protrusions. The critical value of the width below which the elongated protrusions collapse is found in the simulations to be given by a similar force balance which determines the width of the unconfined protrusions, adapted to the flattened geometry (see Supplementary Information). Taken together with our experimental observations, our modeling data suggest that the interplay between the protrusive force of actin polymerization, the intrinsic curvature of the actin nucleating complex, and membrane bending energy could determine the protrusion morphology of cells migrating in confined environments.

### Polarization under confinement relies on processive bleb-based protrusions in the absence of WAVE

In addition to using actin-rich pseudopods for motility, migratory cells build bleb-based protrusions in certain microenvironments (Lämmermann and Germain, 2014; Paluch et al., 2016). These types of protrusions depend on high actomyosin contractility and occur when local weakening of membrane-to-cortex-attachments leads to the separation of the plasma membrane from the underlying actin cortex. Hydrostatic pressure then drives the expansion of this detached segment of membrane to form a protrusion initially lacking filamentous actin (Paluch and Raz, 2013). Since WAVE-null cells are impaired in branched actin assembly at the leading edge, we asked whether they may instead rely on blebbing to generate polarity under confinement. For this purpose, we investigated leading edge dynamics in confined WAVE-null cells expressing an actin-binding fragment of utrophin (Utr261), which associates with cortical actin networks in neutrophils (Fritz-Laylin et al., 2017). Formation of a bleb generally results in a protrusion initially lacking any actin filaments (Bergert et al., 2012); and, consistent with this behavior, confined WAVE-null cells exhibited leading edges with portions of the cytosol extending beyond the boundary of the cortical actin network (Fig. 6A). Intriguingly, the bleb-based protrusions we observed in WAVE-null cells occurred via processive extensions of the plasma membrane that travelled around the cell periphery. At regular intervals, the extension would reverse directions and form a highly-stereotyped serpentine pattern, confining processive extensions to a small section of the plasma membrane to form a leading edge (**Video 15**). This mode of protrusion extension resembles long-known ‘circus movements’, where a bleb forms and progressively extends around the cell in a clockwise or counter-clockwise direction (Loeb, 1928). However, circus movements only occasionally undergo direction reversals (Charras et al., 2008), whereas serpentine protrusions in confined WAVE-null neutrophils did so frequently, on the order of seconds. These frequent reversals appeared to lead to increased directional persistence in protrusion extension, with blebbing confined to a small angular sector of the cell.

**Figure 6.**
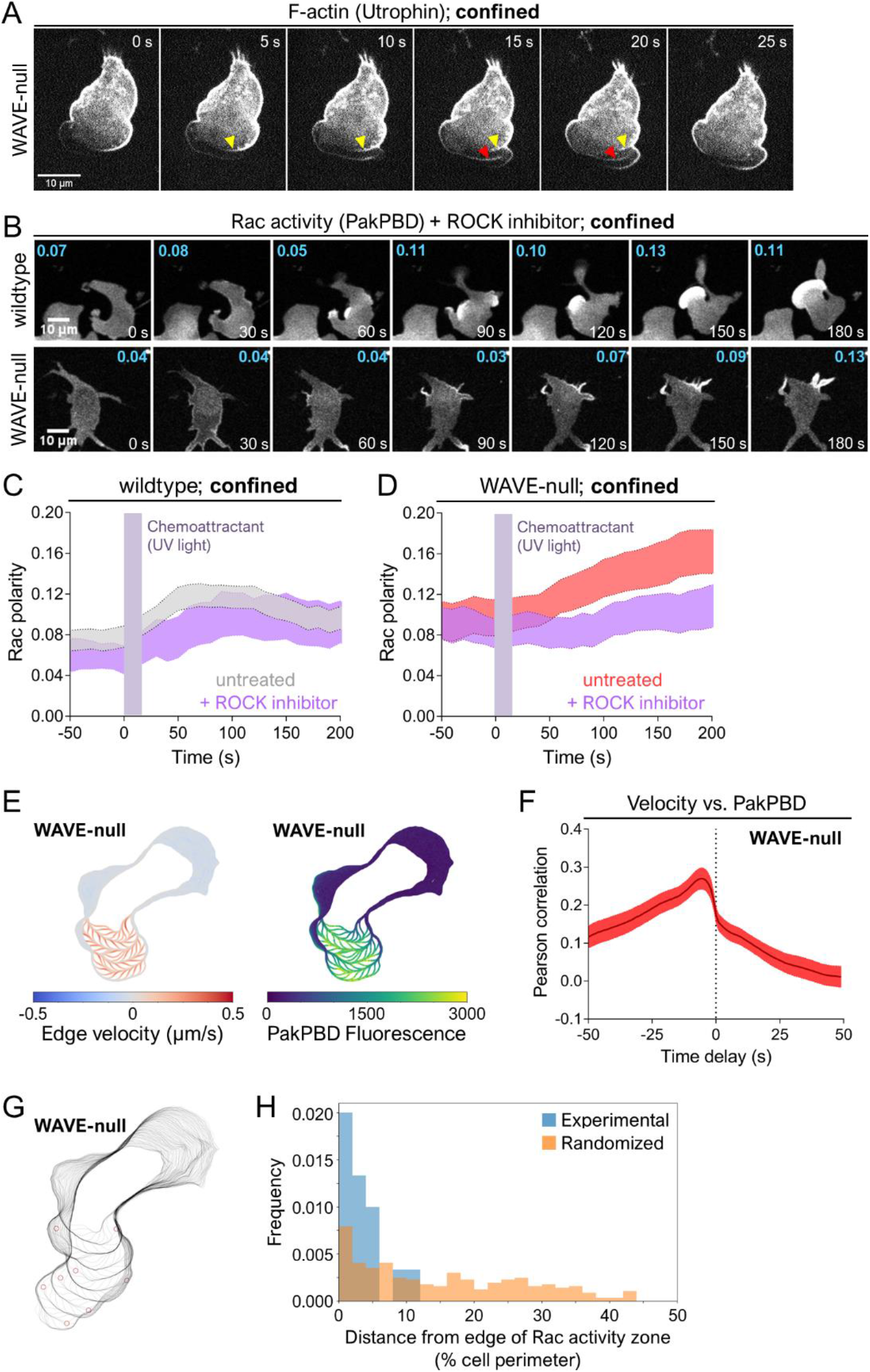
WAVE-null cells under confinement migrate using bleb-like protrusions. **A)** dHL-60 cells expressing the F-actin marker Utr261-mCherry were plated on fibronectin-coated glass in media containing 10 μM caged-fMLP and imaged every 1 s by confocal microscopy. Prior to imaging, cells were confined as in 5A and fMLP was photo-uncaged. Yellow and red arrows indicate where cortical actin networks are left behind following formation of a bleb. **B)** dHL-60 cells expressing the Rac biosensor PakPBD were plated on fibronectin-coated glass in media containing 10 μM caged-fMLP and 20 μM Y27632 and imaged as in 5B. Prior to imaging, cells were confined as in 5A. Values in cyan indicate the degree of Rac polarization, as described in 2C, for the topmost cell fully inside each panel. **C-D)** Quantification of PakPBD polarity as described in 2C for cells prepared as in 6A. Violet bars indicate when UV was used to photo-uncage caged-fMLP (0-20 s). Light shaded regions, +/− 95% CI of the mean Rac polarity. **C)** N = 53 Y27632-treated wildtype cells pooled from 2 independent experiments. Data for “untreated” cells are reproduced from wildtype cells in 5C to aid in comparison. **D)** N = 31 Y27632-treated WAVE-null cells under confinement pooled from 2 independent experiments. Data for “untreated” cells are reproduced from WAVE-null cells in 5C to aid in comparison. **E)** Overlays of extracted cell boundaries over time displaying associated edge velocity (left) and Pak-PBD edge fluorescence (right). **F)** Analysis of Pearson correlation between edge velocity and Pak-PBD fluorescence as a function of temporal offset in fluorescence. The peak Pearson correlation occurs when fluorescence values of Rac activation are shifted back in time by 6 s relative to membrane extension, indicating that membrane extension precedes changes in local Rac activation by 6 s. Red line and shaded area, mean +/− 95% CI. N = 26 cells pooled from 2 independent experiments. **G)** Overlays of extracted cell boundaries as in 6E with red circles marking locations of computationally-identified reversals in bleb propagation direction. **H)** Analysis of spatial coincidence of bleb reversals and boundaries of Rac activity zone. To generate the randomized distribution (orange), the Rac activity zone was rotated randomly and the minimal distances were recalculated. This randomization process was repeated 20 times and the results pooled. Histograms were smoothed with a KDE using Gaussian kernel of bandwidth 20. N = 144 reversals.

As blebbing is powered by actomyosin contractility, we next asked whether the serpentine protrusions of confined WAVE-null cells showed a similar requirement for myosin activity. We treated neutrophils with the ROCK inhibitor Y27632 to block myosin contractility (Maugis et al., 2010). In other contexts, ROCK is required for blebbing but is dispensable for actin protrusion-based chemotaxis (Liu et al., 2015a). Confined wildtype neutrophils treated with Y27632 showed little change in polarized Rac activity, consistent with protrusion growth and cell movement being independent of actomyosin-based contractility in these cells (Figs. 6B-C). In contrast, WAVE-null cells treated with Y27632 under confinement showed reduced polarization of Rac activity upon stimulation and formed only finger-like protrusions (Figs. 6B **and** 6D). However, Y27632-treated WAVE-null cells under confinement still exhibited greater polarity than unconfined WAVE-null cells (Fig. 2D), suggesting that other processes independent of myosin contractility contribute to polarization of Rac activity. Together, our data show that protrusion generation in confined WAVE-null cells arises from actomyosin-dependent blebbing.

### A potential Rac-mediated positive feedback circuit for bleb-based polarization

Established models describing polarization specify the initial production of short-range positive feedback that amplifies small fluctuations in polarity factor activity and long-range negative feedback that acts to limit the spread of the positive feedback and suppress secondary fronts. The combined actions of these two types of feedback lead to the consolidation of polarity to a single site (Meinhardt, 1999). Along with the role of actin polymerization in generating long-range inhibition (Houk et al., 2012; Kozlov and Mogilner, 2007), there is also a well-established requirement for actin assembly in mediating short-range positive feedback during polarization of migratory cells (Huang et al., 2013; Inoue and Meyer, 2008; Nguyen et al., 2016; Sasaki et al., 2007; Wang et al., 2002; Weiner et al., 2007; Yang et al., 2016; Fig. 1A, green arrows). The highly polarized Rac activity we observed in confined WAVE-null cells suggested the presence of short-range positive feedback loops in maintaining localized protrusion growth, despite their impaired branched actin assembly. To explore this possibility further, we imaged cells expressing PakPBD with high temporal frequency to assess whether Rac activity correlated with growth of blebs. Confined WAVE-null cells showed high levels of Rac activity coinciding with the location of serpentine protrusions (i.e. the leading edge) (**Video 16**). To investigate the relation of Rac activity and blebbing more quantitatively, we performed a cross-correlation analysis in which we divided the plasma membrane into 1000 discrete segments and quantified both edge velocity and PakPBD fluorescence for each membrane segment over time (Fig. 6E **and Video 17**; see “Materials and Methods” for details). Edge velocity and PakPBD showed an average instantaneous Pearson correlation coefficient of 0.17 (Fig. 6F). We next shifted the PakPBD intensity measurements for a given timepoint relative to the edge velocity measurements to assess whether local increases in Rac activity came before or after local increases in edge velocity. The correlation between PakPBD fluorescence and edge velocity was highest (Pearson correlation coefficient of 0.27) when PakPBD signal was shifted backwards in time by 6 s relative to edge velocity (Fig. 6F). This analysis indicates that the localized extension of bleb-based protrusions slightly precedes Rac activation and further suggested that Rac activity may restrict the region of the plasma membrane that is permissive for protrusion extension. To test this idea, we computationally identified locations where processive blebbing events underwent reversals (e.g., Fig. 6G and Fig. S4) and determined the distance (as a percentage of total cell perimeter) between reversal location and the edge of the Rac activity zone (see “Materials and Methods” for details). The median distance at which reversals occurred, 3 ± 3%, differed substantially from a distribution where reversals occur at random sites along the cell perimeter, 15 ± 12% (Fig. 6H). These data are consistent with Rac activity providing a permissive zone for bleb propagation, and in conjunction with our observation that blebbing also precedes Rac activation, indicates a bleb-based positive feedback circuit for Rac regulation. These data indicate that neutrophils employ additional feedback mechanisms beyond branched actin assembly to enrich Rac activity at the leading-edge. In future work, the ability of confined WAVE-null cells to migrate exclusively using bleb-based protrusions may serve as a useful platform for dissecting the short-range positive feedback loops underlying bleb-based migration.

## DISCUSSION

Polarization of migratory cells requires coordinated positive and negative feedback loops to restrict protrusion growth to a single site. Actin polymerization is thought to underlie both types of feedback (Diz-Muñoz et al., 2016; Huang et al., 2013; Inoue and Meyer, 2008; Sasaki et al., 2007; Wang et al., 2002; Weiner et al., 2007; Yang et al., 2016), but the roles of different types of actin networks in providing each kind of feedback have not been carefully addressed. Here, we show that branched actin assembly is dispensable for producing short-range positive feedback of Rac activation (Figs. 4E-F **and** 5B-C), whereas the importance of actin in providing long-range negative feedback varies with microenvironment (compare Figs. 2A **and** 3B with Fig. 4B). These observations require a revision of the dominant view that branched actin assembly plays essential roles in both types of feedback. Our work further demonstrates how combined genetic and mechanical perturbations can be used to decouple different migration modes and isolate the components regulating each.

For the long-range inhibition that enables fronts to compete with one another, our data show that this negative feedback can be supported by mechanical deformation of the plasma membrane (Fig. 4). When neutrophils migrate in unconfined environments, WAVE-dependent branched actin assembly is required for generating this force (Figs. 1 **and** 2). However, upon confinement, the resulting membrane extensions arising from this perturbation can compensate for the polymerization force normally provided by branched actin assembly (Fig. 5). This confinement puts WAVE-null cells in a regime where producing either finger-like actin-rich protrusions or bleb-based protrusions can now provide enough protrusive force to satisfy the long-range negative feedback requirement for polarity.

How do cells read out these forces to constrain the levels and spatial distribution of leading-edge regulators? In previous work (Houk et al., 2012), we showed that membrane tension is likely to regulate this long-range inhibitor: Increasing membrane tension suffices to inhibit leading edge signals, whereas decreasing membrane tension prevents their restriction. Similarly, micropipette-based aspiration of plasma membrane at the rear of a migrating keratocyte leads to changes in leading-edge actin dynamics and protrusion velocity within seconds (Mueller et al., 2017). Such data are consistent with locally-applied mechanical forces rapidly propagating through the entire cell. However, it has been difficult to discern whether these mechanical forces depend on membrane tension directly or other correlated physical properties, such as cell shape or local membrane curvature. Recent work has even called into question whether changes in membrane tension can propagate across the cell rapidly enough to serve as a global integrator of mechanical forces (Shi et al., 2018). In other instances, it has been possible to discriminate between the effects of multiple mechanical inputs on cell signaling (e.g. substrate stiffness vs. cell spread area on YAP/TAZ regulation) through the careful design of conditions capable of decoupling these inputs (Dupont et al., 2011). Here we took a similar approach by leveraging a mechanical perturbation (mild hypotonic treatment) which rescued the ability of WAVE-null cells to constrain Rac activity. Importantly, this perturbation does not increase membrane tension despite causing significant changes to cell shape—up to a two-fold increase in volume (Fig. 4D). These results disfavor mechanisms where the mechanosensor restricting Rac activation responds directly to membrane tension, suggesting that tension sensors such as stretch-activated ion channels (e.g., Piezo (Cox et al., 2016; Shi et al., 2018)), cannot be the sole class of mechanosensors mediating neutrophil long-range inhibition. Given that osmotically-induced swelling/shrinking leads to changes in membrane geometry (Cheung et al., 1982; Ting-Beall et al., 1993), mechanosensors responding to changes in membrane shape, using curvature-sensitive membrane binders like BAR proteins (Mesarec et al., 2016; Mim and Unger, 2012), are likely to be involved. An analogous mechanism operates in budding yeast where the highly-conserved mTORC2 signaling pathway is engaged by proteins sensing changes in plasma membrane geometry (Berchtold et al., 2012). Furthermore, we found in neutrophils that mTORC2 signaling restricts leading edge polarity in response to cell stretching (Diz-Muñoz et al., 2016).

Along with generating long-range negative feedback, actin assembly is thought to provide short-range positive feedback to enable polarity (Krause and Gautreau, 2014; Nguyen et al., 2016). However, assessing the role of actin assembly in positive feedback has been challenging. Much of this difficultly has stemmed from a lack of tools for decoupling actin’s role in both types of feedback. Pharmacological inhibitors of actin polymerization like latrunculin target all actin networks and would be expected to break all actin-based feedback (i.e. resulting in pleiotropic effects). Even our more focused approach of targeting only WAVE (Figs. 1-2) or Arp2/3 complex (Fig. 3) initially failed to provide this decoupling: the actin networks assembled by each complex generate mechanical forces at the plasma membrane that are required for long-range negative feedback. It was only when we combined genetic perturbations (i.e. knockout of *HEM1* or *ARP2*) with mechanical perturbations (to generate long-range negative feedback) that we were able to isolate and dissect the role of branched actin assembly in providing short-range positive feedback. This combined approach revealed that branched actin assembly is not required for short-range positive feedback, in contrast to the general assumptions in the field (Iglesias and Devreotes, 2012; Krause and Gautreau, 2014; Saha et al., 2018; Stanley et al., 2014).

When placed under confinement, cells lacking WAVE complex revert from having finger-like protrusions to sheet-like protrusions that resemble wildtype morphology (Figs. 6A-D), a result which is captured even by a highly simplified model describing the interplay between actin-based protrusive force and membrane bending energy (Fig. 5D). Our observations that WAVE-null cells polarize and migrate using blebs are consistent with prior work showing that neutrophils and other migratory cells use distinct strategies to achieve migration in different external environments (Bergert et al., 2012; Liu et al., 2015b; Noselli et al., 2019; Wilson et al., 2013; Yip et al., 2015). However, neutrophils migrating using this bleb-based mode strongly polarize Rac activity to a small section of the plasma membrane (Fig. 5B-C) and Rac activity coincides with protrusion formation and extension (Fig. 6E-F). These results suggest that a branched actin-independent feedback circuit organizes the local positive feedback promoting Rac activity during this mode of migration. It is striking that neutrophils may rely on Rac activity to organize motility around both actin-rich and bleb-based protrusions, despite these protrusion types relying on very different mechanisms for their construction and for leading edge advancement. Other cell types that migrate using blebs, such as zebrafish primordial germ cells (PGCs), rely on Rac activity to support bleb growth (Kardash et al., 2010). Bleb-based motility in PGCs also relies on Cdc42-dependent ‘wrinkling’ of the plasma membrane to create membrane reservoirs that can be released as blebs form and expand (Goudarzi et al., 2017). Since the Cdc42/WASP axis supports neutrophil polarity (Fritz-Laylin et al., 2017; Yang et al., 2016), it would be interesting to test whether this pathway contributes to membrane wrinkling in neutrophils. Such a mechanism would enable neutrophils to extend large amounts of membrane stores when moving through complex microenvironments in vivo or to buffer themselves against changes in cell volume caused by fluctuations in osmolarity. Rac activity may also perform the same function during both migration modes, for example, by promoting local weakening of membrane-to-cortex attachments (MCAs) to facilitate leading extension. In neutrophils migrating using actin-rich protrusions, local depletion of MCAs coincides with leading edge formation (Liu et al., 2015a). Similarly, bleb formation often relies on disruption of MCAs (Charras and Paluch, 2008).

In summary, we used a combination of genetic and mechanical perturbations to show that neutrophils use two mechanistically distinct programs for generating polarity. The first relies on assembly of branched actin networks at the leading edge and is indispensable for polarization in microenvironments where adhesion to substrate is required for motility. However, in microenvironments where neutrophils experience compression forces, a second hidden bleb-based program can maintain polarity even if the branched actin-dependent program is broken. By decoupling changes in membrane tension from changes in cell volume, we further show that polarity-regulating mechanosensors are unlikely to respond directly to tension changes but may instead distinguish changes in membrane morphology, such as local curvature (Gov, 2018). Going forward, leveraging such well-defined environmental perturbations will be helpful in deconvolving the pleiotropic effects arising when disrupting components with multiple cellular functions. This strategy will be particularly important for dissecting processes such as directed migration, where a complex interplay between cell-intrinsic and cell-extrinsic factors underlies cell physiology.

## MATERIALS AND METHODS

### Cell culture

Performed essentially as previously described (Graziano et al., 2017). HL-60 cells were grown in RPMI 1640 media supplemented with L-glutamine and 25 mM HEPES (Corning) and containing 10% (v/v) heat-inactivated fetal bovine serum (Gibco). Cultures were maintained at a density of 0.2-1.0 million cells/mL at 37°C/5% CO_2_. Differentiated HL-60 cells (dHL-60s) were obtained by adding 1.5% (v/v) DMSO (Sigma-Aldrich) to actively growing cells at a density of 0.3 million/mL followed by incubation for an additional 4-5 days. HEK293T cells (used to generate lentivirus for transduction of HL-60 cells) were grown in DMEM (Corning) containing 10% (v/v) heat-inactivated fetal bovine serum (Gibco) and maintained at 37°C/5% CO_2_.

### Plasmids

A vector for mammalian expression of PakPBD-mCitrine was generated by PCR amplification of previously-described PakPBD (Weiner et al., 2007) and mCitrine, and the PCR products were ligated into the pHR backbone between the MluI and NotI sites. Guide RNAs with homology to exon 5 of *NCKAP1L/HEM1* (5’ TGTCACGGATTGAAGATCGG 3’) and exon 1 of *ACTR2* (5’ GGTGTGCGACAACGGCACCG 3’) were cloned into the previously-described LentiGuide-Puro vector (Sanjana et al., 2014), Addgene plasmid #52963. A vector expressing human condon optimized *S. pyrogenes* Cas9-BFP was previously described (Graziano et al., 2017).

### Transduction of HL-60 cells

Performed essentially as previously described (Graziano et al., 2017). HEK293T cells were seeded into 6-well plates and grown until ∼80% confluent. For each well, 1.5 μg pHR vector (containing the appropriate transgene), 0.167 μg vesicular stomatitis virus-G vector, and 1.2 μg cytomegalovirus 8.91 vector were mixed and prepared for transfection using TransIT-293 Transfection Reagent (Mirus Bio) per the manufacturer’s instructions. Following transfection, cells were grown for an additional 3 days, after which virus-containing supernatants were harvested and concentrated ∼40-fold using Lenti-X Concentrator (Clontech) per the manufacturer’s instructions. Concentrated viruses were frozen and stored at −80°C until needed. For all transductions, thawed virus was mixed with ∼0.3 million cells in growth media supplemented with polybrene (8 μg/mL) and incubated overnight. Cells expressing desired transgenes were isolated by culturing in growth media supplemented with puromycin (1 μg/mL) or using fluorescence-activated cell sorting (FACS) as appropriate (FACSAria2 or FACSAria3; BD Biosciences).

### Generation of knockout cell lines using CRISPR/Cas9

Wildtype HL-60 cells were transduced with vectors containing puromycin-selectable guide RNAs (gRNAs) targeting *HEM1/NCKAPL1* or *ARP2/ACTR2*. Following selection, cells were then transduced with an *S. pyrogenes* Cas9 sequence fused to tagBFP. Cells expressing high levels of Cas9-tagBFP were collected using fluorescence-activated cell sorting (FACS), after which a heterogeneous population was obtained, as assessed by immunoblot and by sequencing of the genomic DNA flanking the Cas9 cut site. These cells were then diluted into 96-well plates at a density of ∼1 cell per well to generate clonal lines, which were again verified by genomic DNA sequencing and immunoblot. We verified that candidate clonal lines arose from single cells as previously described (Graziano et al., 2017).

### Preparation of dHL-60s for microscopy

For experiments where cells were observed in unconfined environments, 96-well #1.5 glass-bottom plates (Brooks Life Sciences) were coated with a 100 µL solution of 20 µg/mL porcine fibronectin (prepared from whole blood) and 0.4 mg/mL bovine serum albumin (BSA, endotoxin-free, fatty acid free; A8806, Sigma) dissolved in Dulbecco’s Phosphate Buffered Saline (DPBS; 14190-144, Gibco) and incubated for 30 min at room temperature. The fibronectin solution was then aspirated, and each well was washed twice with 300 µL DPBS. For each well, 0.2-0.4 mL of dHL-60 cells in growth media were pelleted at 200 × g for 5 min, resuspended in 100 µL imaging media (Leibovitz’s L-15 with 0.2% (w/v) BSA, 0.4 mM NaOH, and 1 nM fMLP) and plated. For experiments where PakPBD polarity was quantified (e.g., Fig. 2B-D) or where trajectories were measured (Fig. 1C-D), 5 µM CellTracker Red (C34552, Thermo Fisher) or 1 drop/500 mL NucBlue (R37605, Thermo Fisher), respectively, were added to the imaging media. Cells were then incubated at 37°C/5% CO_2_ for >10 min to permit adherence to the glass, followed by one wash using 100 µL imaging media (containing neither CellTracker Red nor NucBlue dyes).

For experiment where cells were observed in confined environments (i.e. using PDMS-based devices), 25-mm round #1.5 glass coverslips were coated/incubated with a fibronectin/BSA solution (as described in the preceding paragraph), washed once with 1 mL DPBS, and once with 0.4 mL deionized water. Immediately prior to plating cells, coverslips were dried under gaseous N_2_. For each coverslip, 0.2-0.4 mL of dHL-60s in growth media were pelleted at 200 × g for 5 min, resuspended in 20 µL imaging media with 5 µM CellTracker Red, plated, and incubated at 37°C/5% CO_2_ for >10 min to permit adherence to the glass. Cells were then washed once using 50 µL imaging media (without CellTracker Red) and 20 µL imaging media with 10 µM nv-fMLP (chemically caged fMLP, prepared as in (Collins et al., 2015)). PDMS devices (see next section for fabrication) were then placed on top of cells as depicted in Fig. 5A, leftmost panel.

### Microscopy hardware

All imaging experiments with the exception of those depicted in Figs. 1B-D were performed at 30°C on a Nikon Eclipse Ti inverted microscope equipped with a motorized laser total internal reflection fluorescence (TIRF) illumination unit, a Borealis beam conditioning unit (Andor), a CSU-W1 Yokugawa spinning disk (Andor), a 60X PlanApo TIRF 1.49 numerical aperture (NA) objective (Nikon), an iXon Ultra EMCCD camera (Andor), and a laser merge module (LMM5, Spectral Applied Research) equipped with 405, 440, 488, and 561-nm laser lines. All hardware was controlled using Micro-Manager (UCSF). For experiments performed under confinement, the PDMS devices were connected with PE 20 tubing (Braintree Scientific) to a vacuum regulator (IRV1000-01B, SMC Pneumatics) which, in turn, was connected to the building-wide vacuum supply.

Imaging experiments depicted in Figs. 1B-D were performed at 37°C on a Nikon Eclipse Ti inverted microscope equipped with a motorized stage (ASI), a Lamba XL Broad Spectrum Light Source (Sutter), 20x 0.75 NA Plan Apo and 60x 1.4 NA Plan Apo objectives (Nikon), and a Clara interline CCD camera (Andor). All hardware was controlled using Nikon Elements.

### Imaging and analysis

ImageJ (NIH), CellProfiler (Broad Institute), RStudio, Excel (Microsoft), custom Python code, and Prism (Graphpad) were used for all image analysis.

For the displacement/trajectory measurements (e.g., Fig. 1C-D), NucBlue-labeled nuclei were imaged using a 20x objective and DAPI filter cube every 15 s for 10 min by widefield epifluorescence microscopy. The NucBlue channel was used to create a binary mask using Otsu’s method, from which the individual nuclei were segmented. For each nucleus, the center of mass of its binary representation was calculated in each frame. Displacements were calculated by measuring center of each nucleus at each timepoint and calculating the distance from its starting point (i.e. at t = 0 s). Cells were omitted from analysis for the following reasons: i) entering or leaving the field of view during the 10-min observation window, ii) achieving a maximum displacement of < 5 µm during the 10-min observation window (i.e. to remove dead cells and debris), iii) undergoing collisions and/or forming clumps that prevented reliable segmentation of individual nuclei. For the trajectory plots depicted in Fig. 1C, the x-y coordinates of the nuclear positions were normalized to (0, 0) at ‘0 s’. All summary statistics described in the text are mean distance ± SEM.

For the polarity measurements (Figs. 2B-D, 3C-D, 4E-F, 5B-C, 6B-D **and** S3), 488-nm and 561-nm lasers were used to image PakPBD-mCitrine and CellTracker Red, respectively, every 10 s for 5 min by spinning disk confocal microscopy. Focal planes were chosen to be in the lower half of the cell, just above the ventral surface. Cells were stimulated with chemoattractant (fMLP) immediately following the ‘60 s’ timepoint. For experiments performed in 96-well plates (unconfined migration) in isotonic media (e.g., Fig. 2B-D), imaging media containing 0.2% BSA and 200 nM fMLP was added to cells 1:1 to yield a final concentration of 100 nM fMLP. For experiments performed in 96-well plates in hypotonic media (Fig. 4E-F), cells were plated in imaging media as described in “Preparation of dHL-60s for microscopy”. An equal volume of deionized water with 0.2% BSA and 1 nM fMLP was added to cells, followed by incubation for 10 min at 30°C. Afterwards, an equal volume imaging media containing 200 nM fMLP and 0.2% BSA was added to cells, yielding a final concentration of 100 nM. For experiments performed in PDMS devices (confined migration), a 365-nm UV LED flashlight (Americans’ Preferred) was used to photo-uncage nv-fMLP by holding the end of the flashlight 3-5 mm above the top of the PDMS chamber for ∼15 s. Prior to each experiment, the power output of the flashlight was measured using a Slim Photodiode Power Sensor (S130C, Thor Labs). Flashlight batteries were replaced when the power output dropped below ∼70% of the initial reading. Prior to analysis, all images were background-subtracted using a ‘dark’ image acquired by blocking all light from reaching the camera and averaging 100 exposures. The CellTracker Red channel was then used to create binary masks of the cell bodies using Otsu’s method and each cell body was eroded inward by ∼1 µm. Cell bodies were tracked as described in the preceding paragraph to account for cell movement during the experiment. At each timepoint, the cell bodies were used as seeds to identify the PakPBD-mCitrine signal associated with each cell using a previously-described propagation algorithm (Jones et al., 2005). The distance *(d)* between the center of mass of the PakPBD-mCitrine signal (weighted by fluorescence intensity) and the center of mass of a congruent PakPBD binary image (representing the cell’s ‘footprint’) were calculated for each cell at each timepoint. To normalize for differences in cell cross-sectional area, the length *(L)* of the major axis of an ellipse that had the same normalized second central moments as the cellular PakPBD signal was determined. The distance *d* was then divided by 0.5*L*, as depicted in Fig. 2C, to obtain a PakPBD polarity score ranging from 0 to ∼1. In all experiments, polarity was quantified at every 10-s interval (beginning at 50 s prior to stimulation with chemoattractant) with the following exception: In experiments where UV light was used to un-cage nv-fMLP, data from the ‘10 s’ and ‘20 s’ timepoints were always omitted due to excessive background cause by UV light (omitted data indicated by violet bars in Figs. 5C, 6C-D **and** S3C). Cells were omitted from analysis for the following reasons: i) touching the edge of the field of view during the 5-min observation window, ii) undergoing collisions and/or forming clumps that prevented reliable segmentation of individual cell bodies or PakPBD signal, iii) failure to express sufficiently high levels of PakPBD-mCitrine to permit reliable segmentation of cell outlines. All summary statistics described in the text are mean ratio ± SEM.

For edge velocity and PakPBD fluorescence measurements (Fig. 6E-F), images were acquired every 1 s for 5 min. Background-corrected PakPBD images were segmented using a three-step process consisting of Gaussian smoothing, intensity-based thresholding, and distance-transform-based erosion. The threshold and degree of erosion were chosen manually to account for differences in the intensity and polarization of the biosensor in fluorescence images to align the boundary of the binary image with the apparent edge of the cell. To facilitate temporal analysis of edge properties, these boundaries were then fit using a spline interpolation consisting of 1000 evenly-spaced points. The indices of points over time were aligned to minimize the Euclidean distance between point sets in space between consecutive time points. This approach allowed for relatively smooth tracking of points along the cell boundary, making temporal comparisons possible (this approach is conceptually similar to a previously-described alignment strategy (Huang et al., 2013)).

Edge velocity at a particular point *P* at time *t* was tracked by calculating the average of the distance transforms of the binary images at times *t-1* and *t+1* and interpolating the value of the signed distance transform at the coordinates of *P*. Edge fluorescence was tracked by interpolating the value of the background-corrected fluorescence image at the coordinates of *P*. Following alignment of the indices of the points, a temporal comparison of protrusion velocity and background-corrected fluorescence was analyzed by Pearson’s cross-correlation function. Percell correlation functions were then averaged over multiple cells to give Fig. 6F as previously done (Machacek et al., 2009).

To identify bleb turning events, previously derived edge velocity maps were smoothed with a Gaussian filter and protrusions were identified by thresholding the smoothed maps at a threshold value of 0.15 µm/sec. Connected components were then labeled (accounting for the periodic nature of the maps in the position axis) and analyzed on a per protrusion basis. The centers of protrusions were tracked by determining the center of mass of the protrusive region at a given time point (again, accounting for periodicity in the position axis). Reversals were called when the displacement of the center of mass changed direction between consecutive time points. To reduce false positive and negative reversal calls, our analyses were restricted to continuous protrusions that underwent three or more reversals during their lifetime. Next, to analyze the spatiotemporal co-occurrence of identified bleb reversal points and ‘edges’ of the Rac signal, the previously derived fluorescence maps were smoothed and segmented using Otsu’s threshold to binarize the signal. Edge positions were extracted from the binary image using the sobel operator. Finally, these analyses were used together to find the minimum distance between each identified turning event and the edge of the zone of Rac activity at the time of reversal.

### Atomic force microscopy

Custom-made chambers were coated for 30 min with fibronectin (prepared as described in the preceding section) and washed once with DPBS. dHL-60s were plated on each dish in growth media and allowed to adhere for at least 10 min at 37°C. After, cells were washed with RPMI supplemented with 2% FBS and 10 nM fMLP (experiments in Fig. 3A) or RPMI supplemented with 0.2% BSA and 10 nM fMLP (experiments in Figs. 1E **and** 4C).

Experiments in Fig. 3A were performed essentially as previously described (Diz-Muñoz et al., 2016). Olympus BioLevers (k = 60 pN/nm) were calibrated using the thermal noise method and incubated in 2.5 mg/ml Concanavalin A (C5275, Sigma) for 1 h at room temperature. Before the measurements, cantilevers were rinsed in DPBS. Tethers were pulled using a Bruker Catalyst AFM controlled by custom-made LabVIEW software mounted on an inverted Zeiss fluorescent microscope. Approach velocity was set to 1 μm/s, contact force to 100 pN, contact time to 5–10 s and retraction speed to 10 μm/s. After a 10 μm tether was pulled, the cantilever position was held constant until it broke. Tethers that took longer than 15 s to break were omitted from analysis as actin polymerized inside of these longer-lived tethers. Resulting force–time curves were analyzed with the Kerssemakers algorithm (Kerssemakers et al., 2006).

Experiments in Figs. 1E **and** 4C were performed as follows. Olympus BioLevers (k = 60 pN/nm) were calibrated using the thermal noise method and incubated in 2.5 mg/ml Concanavalin A (C5275, Sigma) for 1 h at room temperature. Before the measurements, cantilevers were rinsed in DPBS. Cells were located by brightfield imaging, and the cantilever was positioned at any location over the cell for tether measurement. Cells were not used longer than 1 h for data acquisition. Tethers were pulled using a CellHesion 200 from JPK mounted on an inverted Nikon Ti microscope. Approach velocity was set to 1 μm/s, contact force to 100-300pN, contact time to 5–10 s and retraction speed to 10 μm/s. After a 10 μm tether was pulled, the cantilever position was held constant until it broke. Negative and positive forces relate to the angle the cantilever takes, but the sign is arbitrary. By convention, contacting the cell deflects the cantilever towards positive values. Conversely, when the cantilever is pulled downwards by a membrane tether, the values are negative. For experiments involving hypotonic treatment (Fig. 4C), tether force was first measured in cells plated in RPMI with 0.2% BSA and 10 nM fMLP. An equal volume of deionized water with 0.2% BSA and 10 nM fMLP was then added to the cells and tether force in these same cells was measured over a period of 3-15 min following water addition. Analysis of force-time curves was performed using the JPKSPM Data Processing Software.

### Volume measurements

dHL-60s were pelleted at 180 × g for 5 min and resuspended at a density of ∼20,000 cells/mL in RPMI supplemented with 1 nM fMLP. Volumes were measured using a Beckman Coulter Z2 Coulter Counter with a 100-micron aperture probe. The following settings were used: a calibration factor (Kd) of 59.96, a resolution of 256, a gain of 64, and a current of 0.354 mA. Samples of 0.5 mL were measured for each indicated timepoint. At timepoint ‘0 min’, an equal volume of deionized water with 1nM fMLP was added to the sample, and cell volumes were measured at 1-min intervals for 16 min. For each interval, at least 7,000 cells were measured. A Gaussian function was fitted to each volume distribution to determine the mode volume for the time interval. Analysis was performed using SciDAVis.

### Details of the simulations

The Monte-Carlo simulations of the membrane shape provided in this work follow the scheme described in (Fošnarič et al., 2013; Gompper, 1996). The membrane is described as a triangulated surface, with each node representing either a membrane or a protein domain. In this work a network is composed of 3127 nodes, forming a network of approximately 6250 triangles. Nodes can move and bonds can flip according to the Monte-Carlo (Metropolis) algorithm, within the fixed topology of a sphere and with distance between nodes limited between minimal and 1.7 times larger value. The system is initially thermalized. This gives a thermally fluctuating self-avoiding membrane in steady-state, where protein domains are free to laterally diffuse within the membrane. The energy term due to tension is 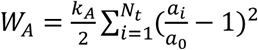 where *k_A_* is the elastic constant of the membrane and the sum runs over all of *N_t_* triangles of the network, *a_i_* is area of triangle *i*, and *a_0_* is area of a tensionless triangle. For *a_0_* we choose an equilateral triangle, 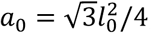 with side lengths *l*_0_ = (*l*_*min*_+ *l*_*max*_)/2. We define membrane tension as the average tension per membrane area, 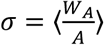, where *A* is the area of the membrane for a given microstate and brackets denote the canonical ensemble average. In addition to the standard bending energy of the membrane 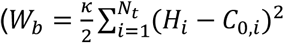, where *κ* is the bending modulus of the membrane, *H_i_*, *C*_*0*,*i*_ the mean and intrinsic curvatures at triangle *i*), there is an energy term that describes the binding between neighboring proteins, with contact energy *w*, as well as an active force of magnitude *F* that acts at every protein node towards the outwards normal. Figs. 1G and S2, show results for a membrane at temperature *T*/*T_0_* = *0.7*, bending modulus *κ* = 20*kT*_0_ with *ρ* = *11%* of active proteins with protrusive force *F* = *1kT_0_*/*l_min_* and direct interaction constant *w = 1kT_0_*. Elastic constants (*k_A_*) of *1kT_0_* and *10kT_0_* are depicted in Figs. 1G and S2, respectively. Fig. 5D shows results for simulations performed as in Fig. 1G with the further addition of two parallel surfaces that confine mobility of the membrane in the ‘z’ direction. Asphericity is defined as previously described (Fošnarič et al., 2013, 2018).

### F-actin staining

Cells were plated in 96-well glass-bottom plates as described in “Preparation of dHL-60s for microscopy” and stimulated using 100 nM fMLP. At indicated timepoints, an equal volume of “fixation buffer” (280 mM KCl, 2 mM MgCl_2_, 4 mM EGTA, 40 mM HEPES, 0.4% bovine serum albumin (BSA), 640 mM sucrose, 7.4% formaldehyde (w/v), pH 7.5) was added to each well and cells were incubated at room temperature for 15-20 min. The fixation solution was then aspirated from each well, cells were washed once with “intracellular buffer” (140 mM KCl, 1 mM MgCl_2_, 2 mM EGTA, 20 mM HEPES, 0.2% BSA, pH 7.5), and 100 µL “staining buffer” (intracellular buffer + 130 nM rhodamine-phalloidin (ThermoFisher) and 0.2% Triton X-100) was added to each well. Following incubation at room temperature for 45 min, the staining buffer was aspirated from each well, cells were washed once with “intracellular buffer” and 100 µL intracellular buffer was added to each well. Cells were stored at 4°C until immediately prior to imaging.

### Fabrication of PDMS devices for cell confinement

PDMS devices were manufactured by mixing Sylgard 184 Silicone Elastomer Base and Sylgard 184 Elastomer Curing Agent (#4019862, Dow Corning) 10:1 (w/w), followed by degassing under vacuum. The exterior suction cup portion of the device was created by placing 3 aluminum rings on a clean silicon wafer in the following order to create concentric circles: i) a disc of 14 mm diameter and 0.5 mm height, ii) a ring of 8 mm inner diameter, 14 mm outer diameter, and 5 mm height, iii) a ring of 19 mm inner diameter, 40 mm outer diameter, and 7 mm height. Degassed PDMS mixture was poured into this mold, a glass slide was used to scape excess off the top, and the device was baked at 80°C for 1 h. Following removal of the device from the metal ring assembly, a 0.75 mm biopsy punch was used to create a hole in the top of the device where vacuum tubing could be inserted.

The second part of the device was fabricated using a silicon wafer with a regularly repeating array of micropillars (440 µm diameter, 5 µm height, spaced 1 mm apart; as previously described (Liu et al., 2015b)). Degassed PDMS mixture was poured onto the wafer, circular #1.5 coverslips (10 mm diameter) were pressed over top of these patterns using tweezers, and these assemblies were baked at 80°C for 1 h. A metal spatula and polyethylene cell lifters (#3008, Costar) were used to remove the PDMS-coated coverslips from the silicon wafer and these coverslips were subsequently attached to the central pillar of the suction cup portion of the device, glass side first. Devices were cleaned by sonication in 70% ethanol for 20 min and placed in a 37°C oven to dry. This cleaning procedure was performed each time a PDMS device was used for an experiment.

### phospho-Pak timecourses

Performed essentially as previously described (Graziano et al., 2017): dHL-60s (4-5 days post-differentiation) were serum-starved by incubation in starvation media (growth media lacking fetal bovine serum) for 45-60 min at 37°C/5% CO_2_ at a density of 1.5×10^6^ cells/mL. For hypotonic treatment experiments (Fig. 4B), cells were then diluted 1:1 using deionized water and incubated for an additional 10 min. Cells were next stimulated with 10 nM fMLP and samples were collected at indicated timepoints by mixing 0.5 mL of cells with 0.5 mL ice-cold stop solution (20% (w/v) trichloroacetic acid (TCA), 40 mM NaF, 20 mM β-glycerophosphate). Samples were incubated at 4°C for 1-12 hours, after which proteins were pelleted, washed once with 0.7 mL ice-cold 0.5% (w/v) TCA, and solubilized in 2x Laemmli sample buffer (Bio-rad). All timecourses were performed at ∼25°C. Samples prepared for validating CRISPR-mediated gene editing (i.e. Fig. S1) were similarly prepared using TCA precipitation.

### Immunoblot assays

Performed essentially as previously described (Graziano et al., 2017): Protein samples in 2x Laemmli sample buffer (prepared from 0.5-1.0 million cells) were subjected to SDS-PAGE, followed by transfer onto nitrocellulose membranes. Membranes were blocked for ∼1 hr in a 1:1 solution of TBS (20 mM Tris, 500 mM NaCl, pH 7.4) and Odyssey Blocking Buffer (LI-COR) followed by overnight incubation at 4 °C with primary antibodies diluted 1:1000 in a solution of 1:1 TBST (TBS + 0.2% w/v Tween 20) and Odyssey Blocking Buffer. Membranes were then washed 3x with TBST and incubated for 45 min at room temperature with secondary antibodies diluted 1:15,000 in Odyssey Blocking Buffer. Membranes were then washed 3x with TBST, 1x with TBS and imaged using an Odyssey Fc (LI-COR). Analysis was performed using Image Studio (LI-COR) and Excel. For phospho-Pak immunoblots, the ratio of phospho-Pak to total Pak was calculated. These values were then normalized by scaling each relative to the value of wildtype cells at timepoint ‘0.5 min’ for cells stimulated with chemoattractant.

Primary antibodies used were Phospho-PAK1 (Ser199/204)/PAK2 (Ser192/197) (Cell Signaling #2605), PAK2 (3B5) (Cell Signaling #4825), WAVE2 (Cell Signaling #3659), Arp2 (GeneTex #GTX103311), and GAPDH Loading Control Antibody (GA1R) (ThermoFisher). Secondary antibodies used were IRDye 680RD Goat anti-Mouse (Li-cor) and IRDye 800CW Goat anti-Rabbit (Li-cor).

## ACKNOWLEDGEMENTS

We thank Alex Groisman, Pooja Suresh, Anne Pipathsouk, and Martin Bergert for experimental assistance, Sean Collins for providing photo-caged chemoattractant, members of the Weiner Lab for helpful discussions, and Kirstin Meyer for a critical reading of the manuscript. Support for this work was provided by a Cancer Research Institute Irvington Postdoctoral Fellowship to B.R.G., the NIH (GM118167 to O.D.W. and T32HL773125 to B.R.G.), the NSF Graduate Research Fellowship Program (Grant No. 1650113 J.P.T. and T.L.N.), the UCSF Moritz-Heyman Discovery Fellowship Program (T.L.N.), the Novo Nordisk Foundation (O.D.W.) and the Center for Cellular Construction (DBI-1548297), an NSF Science and Technology Center. The research work of M.F., S.P., A.I. and V.K.-I. was supported in part by the SI Research Agency (ARRS) grants No. J1-6728, J2-8166, J2-8169, J1-9162, P3-0388 and P2-0232. N.S.G. is the incumbent of the Lee and William Abramowitz Professorial Chair of Biophysics. This study was also supported in part by HDFCCC Laboratory for Cell Analysis Shared Resource Facility through a grant from NIH (P30CA082103).

## SUPPLEMENTAL MATERIALS

**Figure S1.**
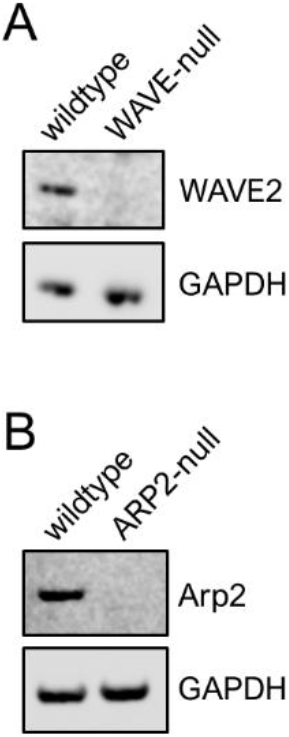
Validation of CRISPR-mediated genome-editing via immunoblot. **A)** WAVE2 antibody immunoblots of wildtype and WAVE-null cells (i.e. lacking Hem-1, the hematopoietic-specific core component of WAVE complex). GAPDH was used as a loading control. **B)** Arp2 antibody immunoblots of wildtype and ARP2-null cells. GAPDH was used as a loading control.

**Figure S2.**
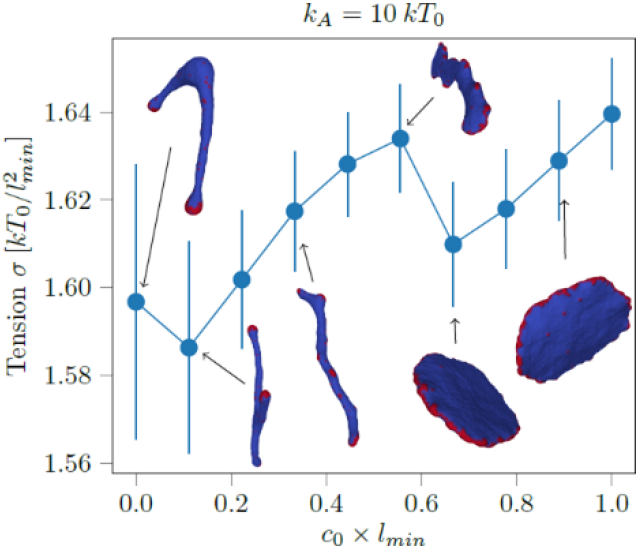
Computational simulations of membrane tension as a function of curvature of actin nucleators. Membrane tension σ (see “Materials and Methods”) as a function of spontaneous curvature, *c_0_*, of actin nucleators. Averaging was performed over 200 statistically independent microstates in equilibrium. Actin nucleators, red. Protein-free bilayer, blue. Error bars, SD. Note that the elastic constant of the membrane (*K_A_*) is 10-fold larger than in simulations performed in 1G.

**Figure S3.**
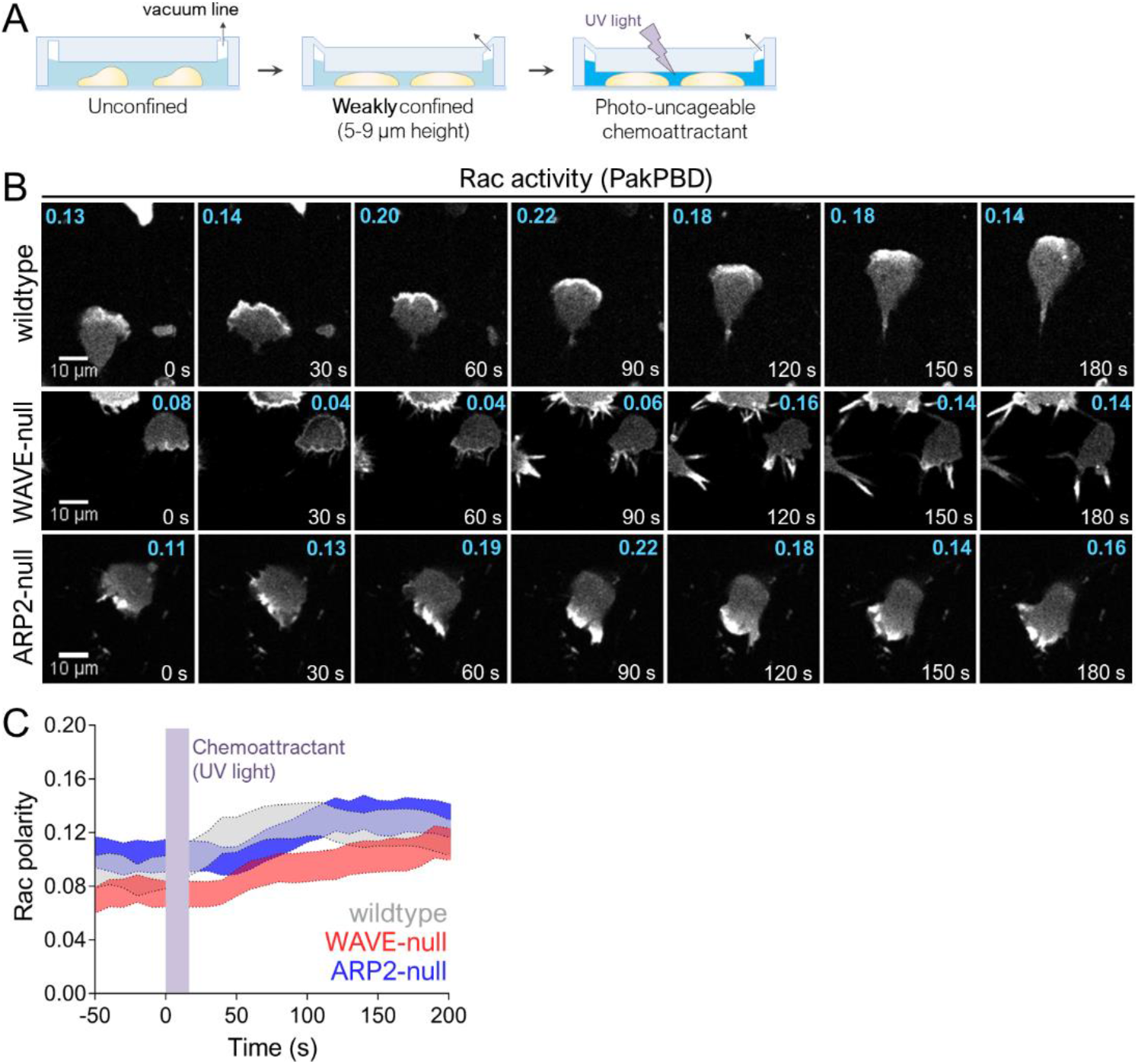
Weak confinement of WAVE-null or ARP2-null cells restores polarized Rac activity. **A)** Schematic for cell confinement experiments. The height of the chamber was set using a vacuum regulator. **B)** dHL-60 cells expressing the Rac biosensor PakPBD were plated on fibronectin-coated glass in media containing 10 μM caged-fMLP and imaged every 10 s by confocal microscopy. Chamber height was set as shown in S3A. Values in cyan indicate the degree of Rac activity polarization, as described in 2C, for the topmost cell fully inside each panel. **C)** Quantification of Rac polarity as described in 2C for cells prepared as in S3B. Violet bar indicates when UV was used to photo-uncage caged-fMLP (0-20 s). Shaded regions, +/− 95% CI of the mean Rac polarity. N = 122 wildtype cells pooled from 3 independent experiments; 143 WAVE-null cells pooled from 3 independent experiments; and 122 ARP2-null cells pooled from 2 independent experiments.

**Figure S4.**
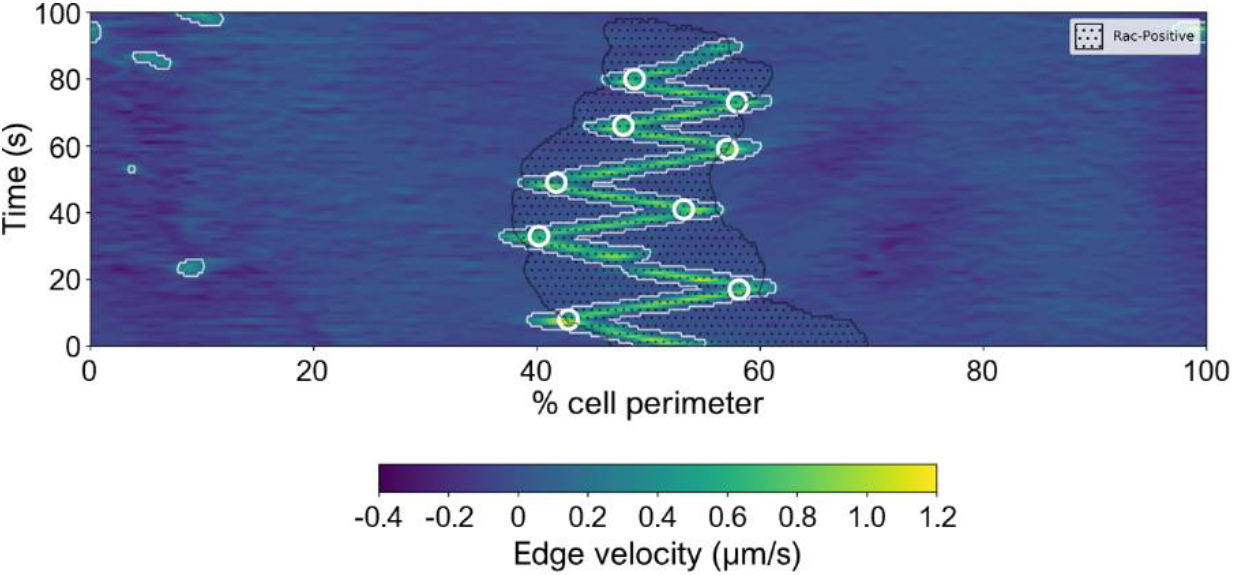
Location of bleb reversals suggests that Rac sets permissive zone for bleb propagation. Kymograph depicting edge velocity map with Rac activity zone overlaid. Computationally-identified reversals are indicated with white circles.

**Video 1.** Wildtype dHL-60s as in Fig. 1B, top panel imaged by DIC microscopy. Time, mm:ss.

**Video 2.** WAVE-null dHL-60s as in Fig. 1B, bottom panel imaged by DIC microscopy. Time, mm:ss.

**Video 3.** Wildtype dHL-60 expressing the Rac activity biosensor PakPBD-mCitrine imaged by confocal microscopy as in Fig. 2B, top panel. Each frame represents a single focal plane.

**Video 4.** WAVE-null dHL-60 expressing the Rac activity biosensor PakPBD-mCitrine imaged by confocal microscopy as in Fig. 2B, bottom panel. Each frame represents a single focal plane.

**Video 5.** Wildtype dHL-60 expressing the Rac activity biosensor PakPBD-mCitrine imaged by confocal microscopy as in Fig. 4F, top panel. Each frame represents a single focal plane.

**Video 6.** WAVE-null dHL-60 expressing the Rac activity biosensor PakPBD-mCitrine imaged by confocal microscopy as in Fig. 4F, bottom panel. Each frame represents a single focal plane.

**Video 7.** Wildtype dHL-60 expressing the Rac activity biosensor PakPBD-mCitrine imaged by confocal microscopy as in Fig. S3B, top panel. Each frame represents a single focal plane.

**Video 8.** Wildtype dHL-60s expressing the Rac activity biosensor PakPBD-mCitrine imaged by confocal microscopy as in Fig. 5B, top panel. Each frame represents a single focal plane.

**Video 9.** WAVE-null dHL-60 expressing the Rac activity biosensor PakPBD-mCitrine imaged by confocal microscopy as in Fig. S3B, middle panel. Each frame represents a single focal plane.

**Video 10.** WAVE-null dHL-60 expressing the Rac activity biosensor PakPBD-mCitrine imaged by confocal microscopy as in Fig. 5B, middle panel. Each frame represents a single focal plane.

**Video 11.** WAVE-null dHL-60s expressing the Rac activity biosensor PakPBD-mCitrine imaged by confocal microscopy. Cells were prepared as described in Fig. 5A. Confinement of cells was repeatedly alternated between a chamber height of 6 µm and 4 µm. Each frame represents a single focal plane.

**Video 12.** Simulation of compression of a vesicle as described in Fig. 5D for *d/l_min_* = 2.0.

**Video 13.** Simulation of compression of a vesicle as described in Fig. 5D for *d/l_min_* = 3.75.

**Video 14.** Simulation of compression of a vesicle as described in Fig. 5D for *d/l_min_* = 7.0.

**Video 15.** WAVE-null dHL-60s expressing the Rac activity biosensor PakPBD-mCitrine imaged by confocal microscopy. Cells were prepared as described in Fig. 5A, except the height of the chamber set to 5 µm. Each frame represents a single focal plane.

**Video 16.** WAVE-null dHL-60s expressing the F-actin marker Utr261-mCherry (Utrophin) imaged by confocal microscopy. Cells were prepared as described in Fig. 5A. Each frame represents a single focal plane.

**Video 17.** WAVE-null dHL-60s expressing the Rac activity biosensor PakPBD-mCitrine imaged by confocal microscopy. Cells were prepared as described in Fig. 5A. Each frame represents a single focal plane.

**Video 18.** WAVE-null dHL-60s expressing the Rac activity biosensor PakPBD-mCitrine imaged by confocal microscopy. Colored lines indicate cell boundaries (see “Materials and methods for details). Images were acquired at 1-s intervals.

## Cell-extrinsic mechanical forces restore neutrophil polarization in the absence of branched actin assembly: Supplementary Information

Before analyzing the shapes of the squeezed vesicles, we can estimate the width of the protrusions of the free vesicle.

### Width of the protrusions of the free vesicle

When the vesicle is in free solution, the conditions that trigger the transition into the “hydra”-shapes (Fig.S1a) are given by the following force balance:

The force applied at the tip of the cylindrical protrusion by the cluster of active proteins is

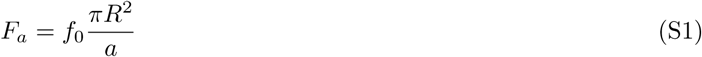

where *f*_0_ is the force per active protein, *R* is the radius of the cylinder, and *a* is the area of a protein on the membrane.

This is balanced by the restoring force of the membrane bending energy

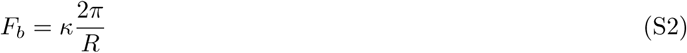

with *κ* the bending modulus. The force balance gives the radius of the cylindrical protrusions in this phase of the vesicle shapes

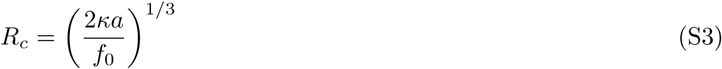

This relation, and especially the predicted scaling of the protrusion radius as (*κ/f*_0_)^1/3^, was verified by simulations (Fošnarič et al., 2018). We next follow a similar calculation to estimate the critical width below which individual protrusions merge to form a continuous cluster along the rim of the confined vesicle.

### Protrusions of a squeezed vesicle

When the cylindrical protrusions are now squeezed to a very narrow, and constant height *d*, it is more natural to define the lateral width of the protrusion *L* (Fig.S1b). The active force is still given by the number of active proteins at the front of the flat protrusion

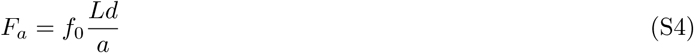

and the bending force

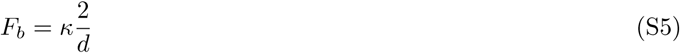

The force balance gives the stable lateral width of these flat protrusions

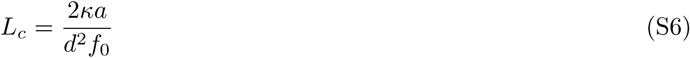

The critical width at which the width of the protrusions merge to form a continuous cluster of proteins along the rim of the squeezed vesicle, can be estimated by substituting in Eq.S6 *L_c_ ∼ R*_vesicle_. This gives us the following scaling between the critical width and the system parameters

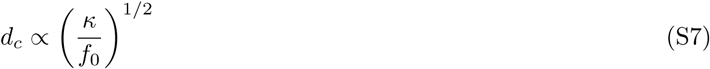

Note that due to the two-dimensional geometry, the scaling is different from that of the width of the free protrusions (Eq.S3).

We tested this scaling using simulations: Since for (*κ/f*_0_) = 20*l_min_* we find that the critical width is *d_c_* ∼ 3.5*l_min_* (Fig.5D), according to the scaling of Eq.S7 we expect the critical width to be:

- *d_c_* ∼ 1.24*_min_* for (*κ/f*_0_) = 2.5*l_min_*. Indeed in simulations we found that the individual protrusions did not merge even when we squeezed down to *d* = 2.25*l_min_*. Below this, the discrete nature of the triangulated surface does not allow for an accurate description of the highly curved rim of the vesicle.
- *d_c_* ∼ 7*_min_* for (*κ/f*_0_) = 80*l_min_*. Indeed in simulations we found the protrusions to form a continuous cluster at the rim of a uniform circular vesicle for all *d* < ∼ 7*l_min_*. For *d* = 8*l_min_* we find that the proteins at the rim clearly form several disconnected clusters.

**FIG. S1.**
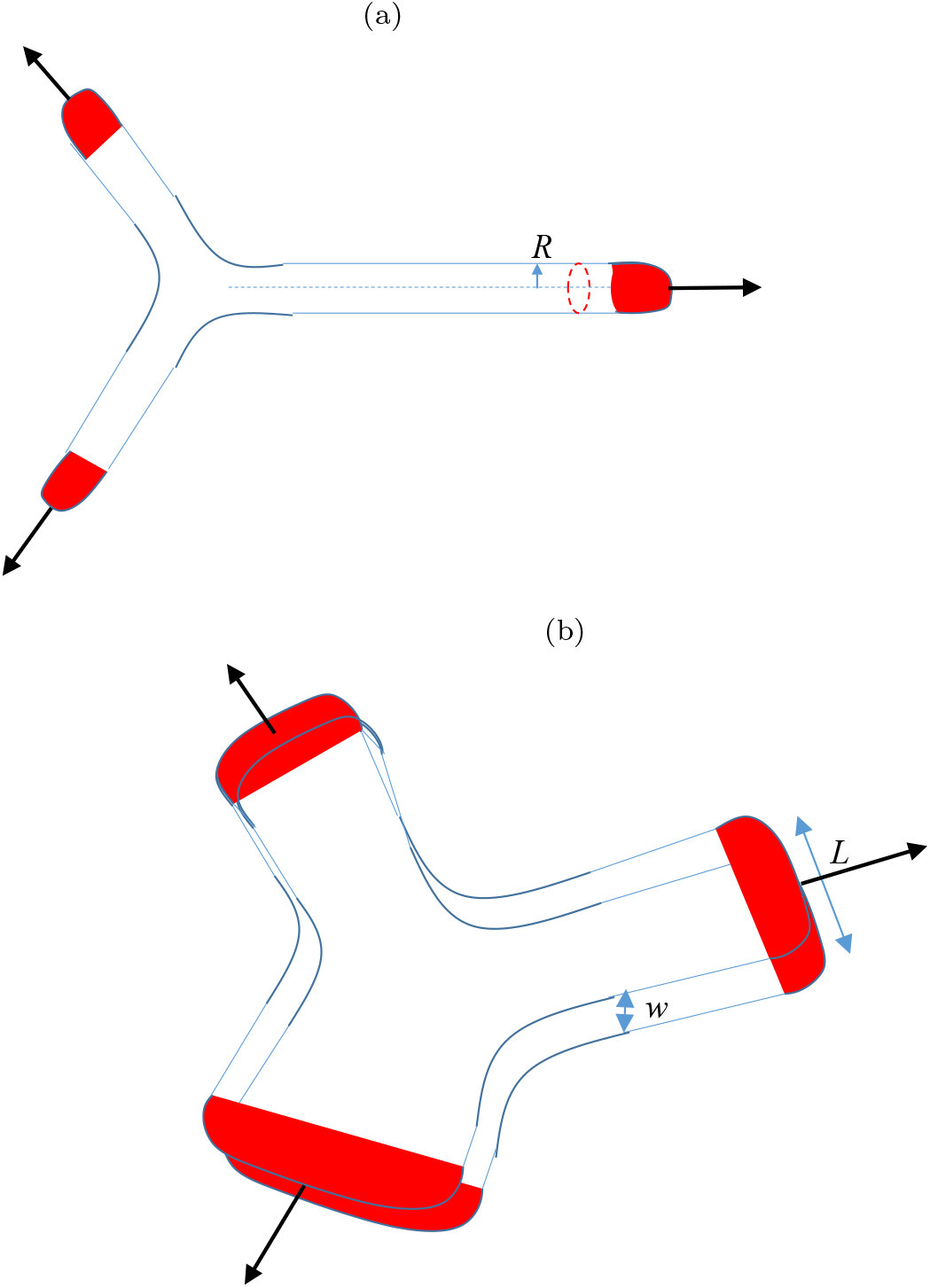
The proteins are denoted by red, and their active force *F_a_* by the black arrow. (a) Cylindrical protrusions of a free vesicle, (b) Flattened protrusions of a squeezed vesicle.

